# Neural stem and progenitor cells support and protect adult hippocampal function via vascular endothelial growth factor secretion

**DOI:** 10.1101/2023.04.24.537801

**Authors:** Jiyeon K. Denninger, Lisa N. Miller, Ashley E. Walters, Manal Hosawi, Gwendolyn Sebring, Joshua D. Rieskamp, Tianli Ding, Raina Rindani, Kelly S. Chen, Sakthi Senthilvelan, Abigail Volk, Fangli Zhao, Candice Askwith, Elizabeth D. Kirby

## Abstract

Adult neural stem and progenitor cells (NSPCs) reside in the dentate gyrus (DG) of the hippocampus throughout the lifespan of most mammalian species. In addition to generating new neurons, NSPCs may alter their niche via secretion of growth factors and cytokines. We recently showed that adult DG NSPCs secrete vascular endothelial growth factor (VEGF), which is critical for maintaining adult neurogenesis. Here, we asked whether NSPC-derived VEGF alters hippocampal function independent of adult neurogenesis. We found that loss of NSPC-derived VEGF acutely impaired hippocampal memory, caused neuronal hyperexcitability and exacerbated excitotoxic injury. We also found that NSPCs generate substantial proportions of total DG VEGF and VEGF disperses broadly throughout the DG, both of which help explain how this anatomically-restricted cell population could modulate function broadly. These findings suggest that NSPCs actively support and protect DG function via secreted VEGF, thereby providing a non-neurogenic functional dimension to endogenous NSPCs.

## Introduction

The dentate gyrus (DG) of the hippocampus is one of only a few brain regions that supports resident neural stem and progenitor cells (NSPCs) in adult mammals^1, 2^. Adult hippocampal NSPCs have generated clinical interest because hippocampus-mediated memory function declines with many injuries and diseases, such as seizures, trauma, and Alzheimer’s disease^3^. Previous research on the functional properties and therapeutic applications of adult NSPCs has focused on their ability to create new neurons that exhibit unique electrophysiological properties^4, 5^. NSPCs, in contrast, are not electrophysiologically active. However, preclinical studies have increasingly identified the growth factors and cytokines secreted by stem cells (their secretome) as a major component of the therapeutic effect of stem cell transplants^6, 7^. The role that the endogenous NSPC secretome plays in brain health remains more sparsely investigated^3, 8, 9^. Nonetheless, several recent studies suggest that NSPCs can shape adult niche properties through their secretome^10–12^.

We recently showed that a major component of the adult DG NSPC secretome is the pleiotropic, soluble protein vascular endothelial growth factor (VEGF, or VEGFA)^13^. We showed that self-derived VEGF signaling in radial glia-like neural stem cells (NSCs) is necessary to maintain their quiescence and prevent premature exhaustion^13^. While this work revealed an autocrine role for NSPC-derived VEGF in maintaining neurogenesis, the potential VEGF paracrine effects were unclear. VEGF is a multi-faceted molecule that can modulate multiple cellular processes in the adult CNS and may protect or impair neuronal function depending on the context^14, 15^. In contrast to the weeks-long process of neurogenic exhaustion, VEGF paracrine signaling has the potential for rapid, acute regulation of the DG. In this work, we therefore investigated the acute, paracrine role of VEGF derived specifically from NSPCs in the function of the adult mouse DG.

## Results

### Knockdown of VEGF in NSPCs impairs hippocampal spatial memory

Our previous research showed that astrocytes and NSPCs are the primary producers of VEGF in the adult DG with NSPCs contributing ∼1/3 of total DG VEGF^13^. To investigate the functional effect of VEGF derived specifically from NSPCs, we crossed NestinCreER^T2^ mice^16^ with VEGF^lox^ mice^17^ (**Fig. 1A**). NestinCreER^T2^ mice drive LoxP recombination with high selectivity in NSPCs when exposed to the synthetic estrogen tamoxifen (TAM)^16, 18, 19^. We confirmed this selectivity of recombination in NestinCreER^T2^ mice crossed with stop-floxed EYFP reporter mice, finding that 97% of EYFP+ cells in the DG were NSPCs and less than 1% were astrocytes 3d after TAM (**Supplementary Fig. 1A-D**). We further confirmed knockdown specificity in adult NestinCreER^T2^;VEGF^lox/lox^ mice (iKD) and their VEGF^lox/lox^ littermates (Wt). After TAM injection, in situ hybridization with a probe against *vegfa* coupled with immunolabeling for GFAP confirmed *vegfa* loss selectively in GFAP+ radial glia-like putative NSCs but not in GFAP+ stellate putative astrocytes (**Fig. 1B,C; Supplementary Fig. 1E**).

**Figure 1.**
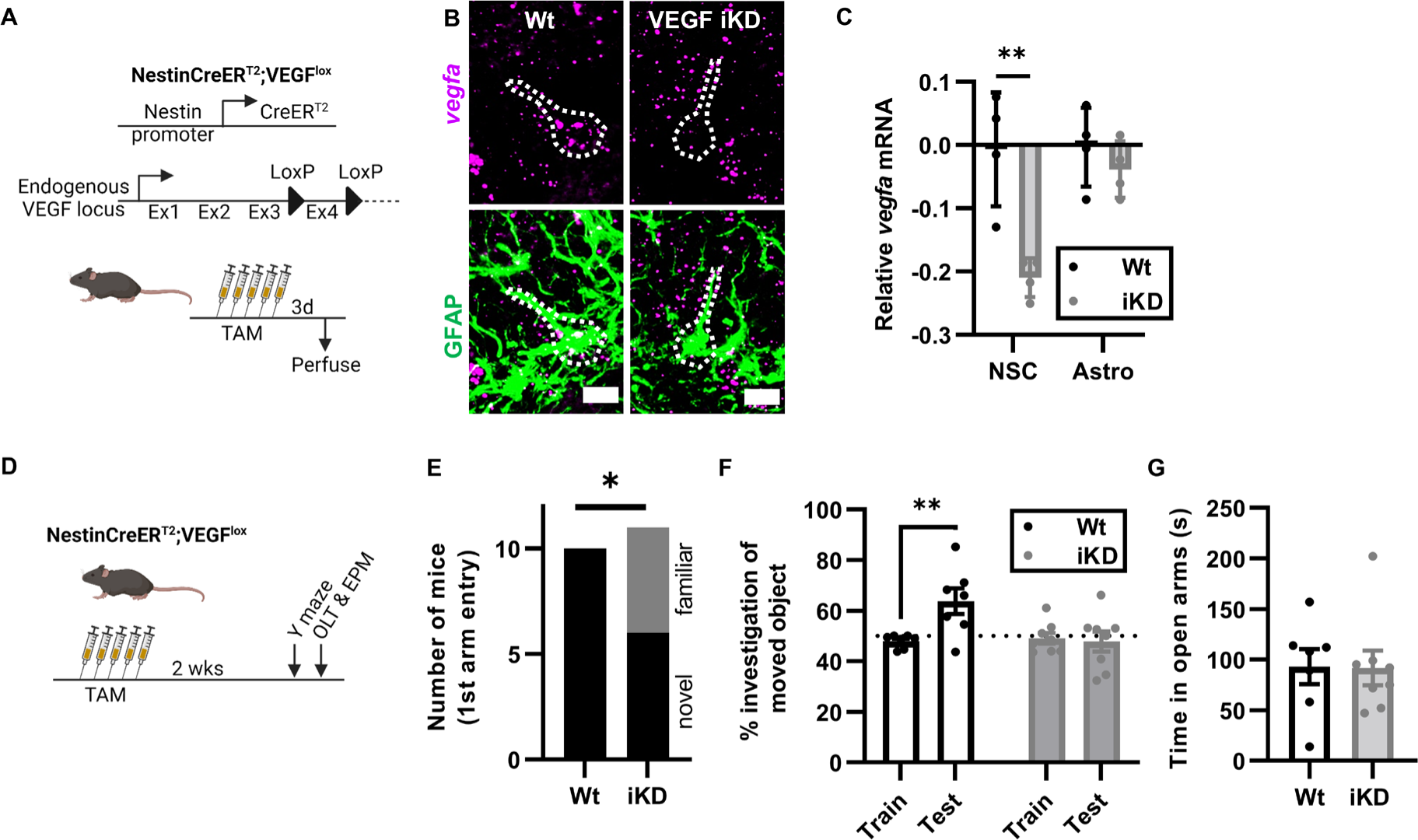
NSPC-VEGF knockdown impairs hippocampal spatial memory. A) Schematic of transgenic mouse strains used. NSPC-VEGF knockdown (İKD) was induced by 5d TAM injection **in** NestinCreER^T2^;VEGF^l0X/l0X^ mice. VEGF^lθx/l0×^ (Crē) littermates were controls (Wt). 3d after TAM, mice were perfused for RNAscope in situ hybridization. B) Representative immunofluorescent images of GFAP immunolabeling with *vegfa* mRNA. Dashed line outlines putative NSC. Scale = 10 µm. C) Average *vegfa* mRNA puncta per cell (log_2_fold change normalized to Wt NSC levels) in Wt and İKD NSCs versus astrocytes. 2 way repeated measures ANOVA cell type x genotype interaction p = 0.0178, genotype p = 0.0021, cell type p = 0.0144. **p = 0.0010 Sidak’s multiple comparisons. Mean ± SEM and individual mice shown. N = 4 Wt, 4 İKD mice. D) Schematic of treatments and behavioral testing. Mice received 5d TAM then 2 weeks later were tested on a hippocampal dependent Y maze, hippocampal dependent object location task (OTL) and an elevated plus maze (EPM). E) Number of mice who chose the novel arm first for entry in Y maze. *p = 0.0351 Fisher’s exact test. N = 10 Wt, 11 İKD mice. F) Percent time spent investigating a moved object in the OLT during training and testing. 2-way ANOVA trial x genotype interaction p = 0.0079. **p = 0.0031 Sidak’s multiple comparisons. Mean ± SEM and individual mice shown. N = 7 Wt, 8 İKD mice. G) Time spent in open arms of the EPM. T-test, ns. Mean ± SEM and individual mice shown. N = 7 Wt, 8 İKD mice.

To test the role of VEGF in hippocampal behavioral functions, we treated VEGF iKD mice and Wt littermates with TAM then assessed performance on hippocampus-dependent spatial-contextual memory Y-maze and object location tasks two weeks later (**Fig. 1D**). We have previously shown that loss of NSPC-VEGF impairs NSC maintenance and thereby neurogenesis, but the loss of new neurons requires > 3 weeks to emerge^13^. In addition, newborn neurons require 3-4 weeks to integrate into local circuitry^20^. This 2-week timepoint for behavioral testing was therefore chosen to be sufficiently close to VEGF knockdown that any changes in production of new neurons would not likely have functional effects on hippocampal circuitry and behavioral changes could be attributed to acute effects of VEGF protein on existing cells.

In a hippocampus-dependent Y maze task, all Wt mice (100%, 10/10) chose to enter a novel, unexplored arm first over a familiar arm, while significantly fewer (54.5%, 6/11) VEGF iKD mice chose the novel arm first (**Fig. 1E**). Latency to enter the novel arm was longer in VEGF iKD than Wt mice but that measure did not reach statistical significance, nor did time in the novel arm (**Supplementary Fig. 1F,G**). In the novel object location task, both Wt and iKD mice showed no preference for either of 2 objects during initial exploration (train). During testing, when one object was moved, Wt mice showed a significant increase in investigation preference for the moved object, but VEGF iKD did not (**Fig. 1F**). There were no differences in total object investigation time, suggesting similar motor ability and motivation to investigate in both groups (**Supplementary Fig. 1H**). Last, we tested mice in an elevated plus maze to assess anxiety-like behaviors^21^. Wt and iKD mice showed no difference in time in the open arms or entries into the closed arms (**Fig. 1G, Supplementary Fig. 1I,J**).

To further confirm these findings, we repeated these tests in a separate cohort of mice 3 days after TAM, reasoning that deficits due to acute loss of VEGF protein could be evident soon after recombination, given the generally short half-life of secreted proteins in vivo (**Supplementary Fig. 1K**). Y-maze first arm choice was not different between Wt and iKD mice, though this analysis was complicated by the failure of the Wt mice to show a preference for the novel arm (**Supplementary Fig. 1L-N**). In the novel object location test, in contrast, Wt mice showed significant preference for the moved object while the iKD mice did not, replicating our finding at 2 weeks (**Supplementary Fig. 1O,P**). In the EPM, iKD mice showed a small but significant increase in time in the open arms, but there was no difference in the time in open arms, compared to Wt mice (**Supplementary Fig. 1Q,R**). No difference in closed arms entries was found (**Supplementary Fig. 1S**). Overall, findings in both the Y maze and object location task suggest that NSPC-derived VEGF acutely supports hippocampal spatial memory, while general object exploration and elevated plus maze findings further suggest that these effects are likely not confounded by any changes in anxiety-like behavior.

### Knockdown of VEGF in NSPCs causes DG hyperexcitation

We next sought to clarify the circuit mechanism by which NSPC-VEGF supports memory function. VEGF is known to signal via VEGFR2 expressed by neurons, generally in support of memory function^22, 23^. In many brain regions, enhanced memory is linked to enhanced neuronal activity. The DG, however, is heavily inhibited by local interneurons, and this inhibition is hypothesized to underpin its role in spatial memory pattern separation^24–27^. We therefore investigated the hypothesis that NSPC-derived VEGF supports hippocampal memory via suppression of DG excitability using slice electrophysiology. Ex vivo hippocampal slices were prepared from Wt and iKD mice 3d after the last TAM injection (**Fig. 2A**). Perforant path stimulation revealed a significant increase in I-V slope in the DG of VEGF iKD mice compared to Wt littermates, suggesting increased basal synaptic transmission in iKD mice DGs (**Fig. 2B**). LTP induced by high frequency stimulation (HFS) of the perforant path was also significantly higher in VEGF iKD mice DGs than Wt littermate controls (**Fig. 2C**). As picrotoxin is present within the bathing solution, these results suggest greater plasticity in excitatory circuits within the DG. No difference was found in paired-pulse ratio (**Fig. 2D**) indicating that the effects observed are likely due to postsynaptic, rather than presynaptic, changes. Together, these data suggest greater excitatory transmission and plasticity following loss of NSPC-derived VEGF supporting the idea that there is hyperexcitability within the DG.

**Figure 2.**
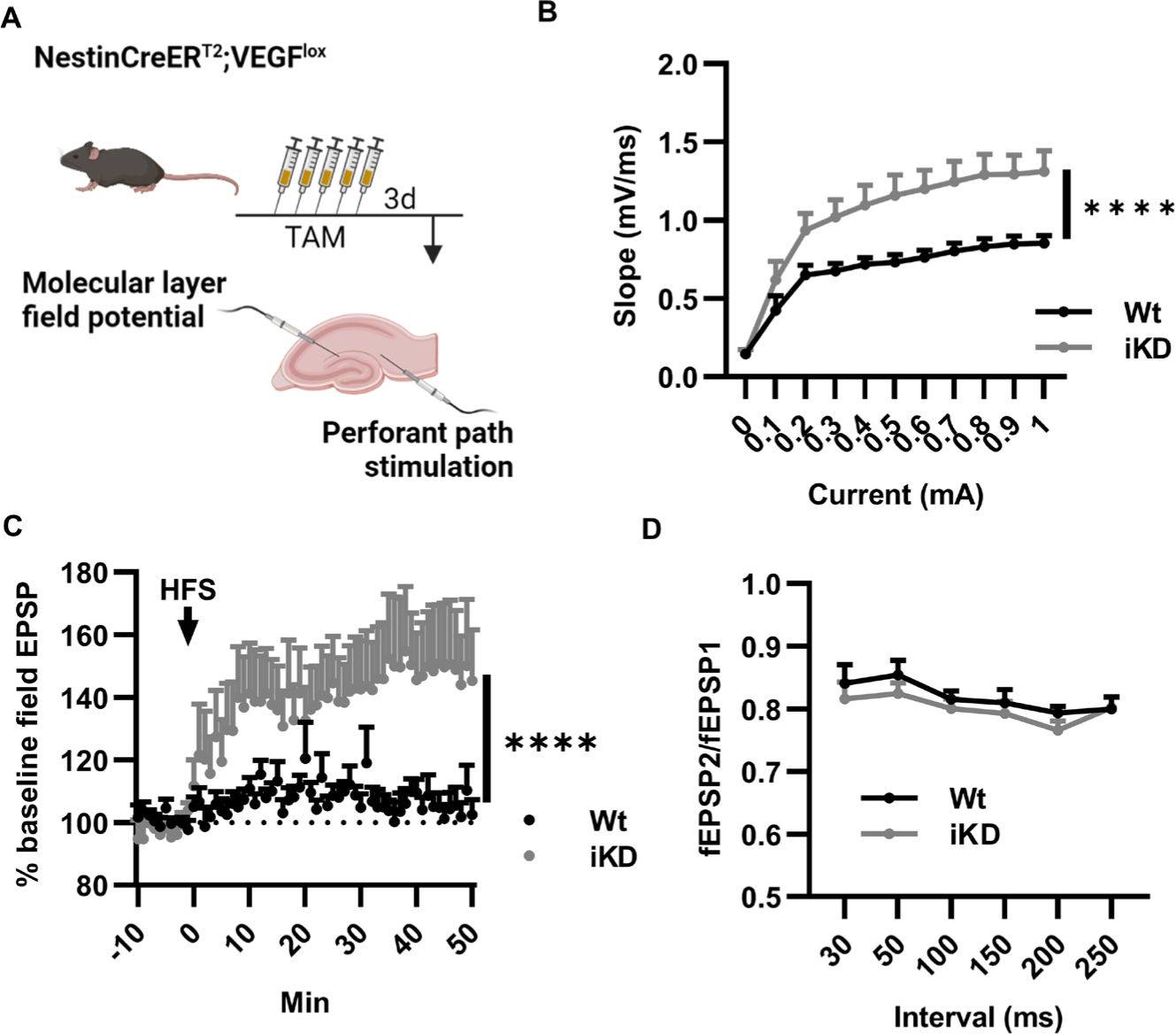
NSPC-VEGF knockdown increases DG excitability. A) Schematic of treatment before ex vivo extracellular electrophysiology with stimulation in the perforant path and recording in the DG molecular layer. B) Slope/current curves. Mean ± SEM shown. 2 way repeated measures ANOVA *”*mA × genotype p < 0.0001, genotype p = 0.0069, mA p < 0.0001. N = 10 Wt, 10 İKD mice, 4-5 slices averaged per mouse. C) fEPSP relative to baseline before and after high frequency stimulation (HFS) to induce LTP. 2 way repeated measures ANOVA ****time x genotype p < 0.0001, time p = 0.0054, genotype p = 0.0491. N = 10 Wt, 10 İKD mice, 4-5 slices averaged per mouse. D) fEPSP2/fEPSP1 in extracellular ex vivo electrophysiological recordings. 2 way repeated measures ANOVA, interval p = 0.0017. N = 10 Wt, 10 İKD mice, 4-5 slices averaged per mouse.

### VEGF disperses widely throughout the DG from a point source

In our previous work^13^, we showed that VEGF in the DG is synthesized primarily by astrocytes and NSPCs. In the adult DG, these two cell types are concentrated in separate DG cell layers, with NSPCs residing exclusively in the subgranular zone and astrocytes residing primarily in the more acellular spaces of the DG like the hilus and molecular layer. VEGF is a secreted, soluble protein, and its potential to regulate other cells will depend critically on its spatiotemporal spread through tissue. Computational estimates suggest that VEGF decreases by up to 12% every 10 µm away from its source in mature muscle tissue^28^. Given that DG subregions can span 50-200+ µm, such a rapid decay in VEGF through tissue would severely limit what cell types NSPC-derived VEGF could reach. We therefore sought to quantify dispersal of VEGF through live adult mouse DG to determine if it is plausible that NSPC-derived VEGF could regulate cells throughout the DG subregions.

To visualize VEGF dispersal through live DG tissue, we used genetic code expansion coupled with bioorthongonal non-canonical amino acid tagging to create tagged forms of both major VEGF isoforms synthesized by NSPCs and astrocytes: VEGF_120_^TAG^ and VEGF_164_^TAG^. We chose this approach over a more traditional approach, such as generating a fluorophore-fusion protein, because most fluorophores (e.g. GFP, 27 kDa) would more than double the molecular weight of VEGF (∼17-22 kDa). This tagging approach, in contrast, involves integrating a single non-canonical amino acid in a recombinant VEGF protein, which can be detected by spontaneous and highly specific inverse–demand Diels–Alder cycloaddition click reaction with tetrazines^29–32^. In brief, we designed expression constructs for each isoform of VEGF cDNA with an in-frame amber stop codon in exon 2. We generated tagged VEGF isoforms using these constructs coupled with expression of a mutant pyrrolysyl-tRNA synthetase/tRNA^Pyl^ pair (PylRS/tRNA_pyl_) that mediates incorporation of a noncanonical amino acid (TCO*A) at amber stop codons^29^ (**Fig. 3A**). Isolated proteins were found at the appropriate weight for VEGF_120_ and VEGF_164_ dimers respectively (∼34 and ∼44 kDa) using 3 different methods: 1) at the total protein level, 2) reacted with streptavidin IR800 (to reveal TCO*A-tetrazine-biotin presence) and 3) using VEGF immunoreactivity (**Supplementary Fig. 2A**). Tagged VEGF monomers were also found at the predicted weight when separated under reducing conditions (∼17 and ∼22 kDa) (**Supplementary Fig. 2B**).

**Figure 3.**
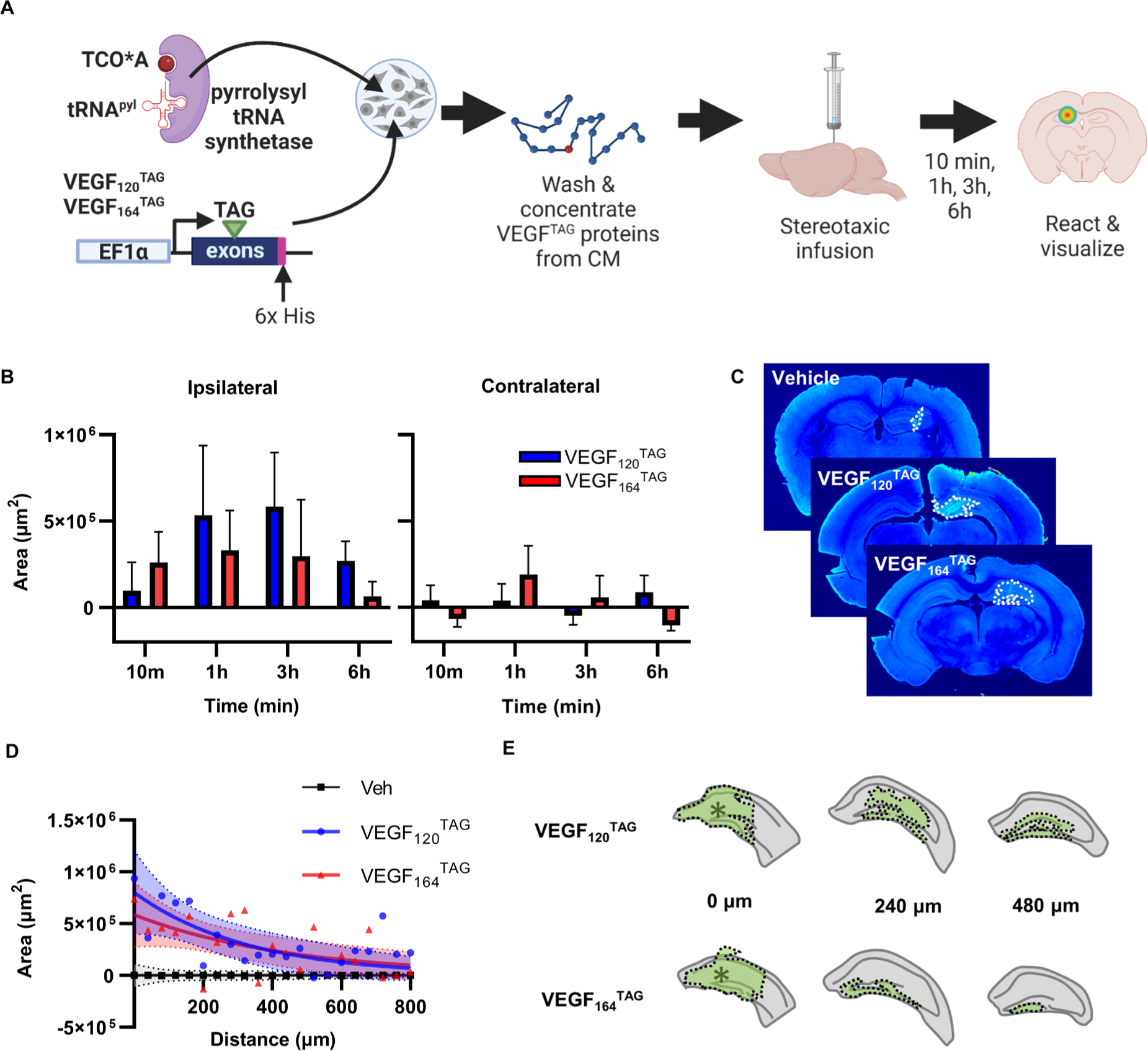
VEGF disperses widely from a point source in the DG. A) Schematic of experimental design. VEGF^TAG^ constructs for 120 and 164 isoforms were generated with an amber stop codon (TAG) in exon 2. VEGF^TAG^ constructs were transiently expressed in HEK cells, along with a pyrrolysyl tRNA synthetase/tRNApy’ pair. TCO*A was provided to generate tagged VEGF proteins via amber stop codon suppression. VEGF proteins were isolated, click reacted with tetrazine-biotin and concentrated from conditioned media of transfected cells then stereotaxically infused in the DG of adult wildtype mice. Mice were perfused 10min-6h later and TCO*A-bearing proteins were detected using infrared imaging. B) Area of VEGF^TAG^ detection in DG ipsilateral and contralateral to the infusion at the epicenter of infusion over time (minus vehicle as baseline). 3 way ANOVA hemisphere p = 0.0014 all other ns. Mean shown as points with SEM and shaded area. N = 4-5 mice per isoform per timepoint. C) Representative images of VEGF^TAG^ areas at the injection epicenter in a vehicle mouse, a VEGF_120_^tag^ (3h) mouse and a VEGF_164_^TAG^ (1 h) mouse. D) Area of VEGF^TAG^ detection in DG through the DG from the epicenter (0 µm). Mean shown as points. One phase decay non-linear fit as line and 95% confidence intervals as shaded area. VEGF_120_^TAG^: Area = 801969*exp(−0.003044*distance). VEGF_164_^TAG^: Area = 586163*exp(−0.002220*distance). Veh: Area = 0.05969*exp(−0.004177*distance). N = 4 mice per isoform; N = 9 vehicle mice. E) Traces of representative VEGF^TAG^ areas in single mice from epicenter to −480 µm rostral. Dashed outline filled green shows VEGF^TAG^+ area, ‘indicates point of needle infusion.

Adult wildtype mice were unilaterally stereotaxically infused with VEGF_120_^TAG^, VEGF_164_^TAG^ or the vehicle control into DG then perfused 10 min, 1hr, 3h or 6h later. The dispersion area of VEGF^TAG^-bearing proteins was quantified using automated region detection software. At the epicenter of the infusion, we found that both VEGF isoforms reached maximal dispersion ipsilateral to the injection site 1-3h after infusion, with minimal spread to the contralateral DG (**Fig. 3B,C; Supplementary Fig. 2C**). There was no significant difference between VEGF isoforms, though VEGF_164_^TAG^ appeared to have slightly less dispersal area than VEGF_120_^TAG^ at later timepoints, likely reflecting the known lower solubility of VEGF_164_ compared to VEGF_120_. To quantify the spatial dispersal of VEGF^TAG^ though the DG, we focused on the decay of signal over space from the epicenter. We found that the decay half-life of spatial dispersal was similar between isoforms: 227.7 µm (95% CI 95.94 to 906.3 μm) for VEGF_120_^TAG^ and 312.2 µm (95% CI 131.8 to 1612 μm) for VEGF_164_^TAG^ (**Fig. 3D**). Representative areas show that both isoforms largely fill the area of the DG within that half-life (**Fig. 3E, Supplementary Fig. 2D**). These findings suggest that VEGF in the adult DG has substantial spatial dispersal from a point source. The location of NSPCs within the SGZ is therefore unlikely to be a barrier to wide-spread signaling of their secreted VEGF.

### DG VEGF derives from astrocytes and NSPCs following excitotoxicity

Several studies suggest that VEGF may help prevent DG hyperexcitation and neuronal damage in pathological conditions^33–35^. Our findings that loss of NSPC-VEGF increases DG excitability and that VEGF likely can disperse widely throughout the DG from isolated sources suggests a possible role for NSPCs in reducing widespread pathological excitotoxicity, such as that which occurs with seizure-related activity. We therefore next investigated the hypothesis that VEGF protects the adult mouse DG from seizure-related excitotoxic damage.

To determine the relative contribution of NSPCs to total DG VEGF after seizure-induced excitotoxic injury, we used the glutamatergic agonist kainic acid (KA). First, we treated VEGF-GFP transcriptional reporter mice with KA (or vehicle) and quantified GFP expressing astrocytes, NSCs and intermediate progenitor cells (IPCs) 1, 3 and 7d later. The density of VEGF-GFP+ astrocytes and NSCs increased with KA, peaking around 3-7d after injury, in parallel with an increase in total astrocytes and NSCs (**Fig. 4A,B; Supplementary Fig. 3A,B**).

**Figure 4.**
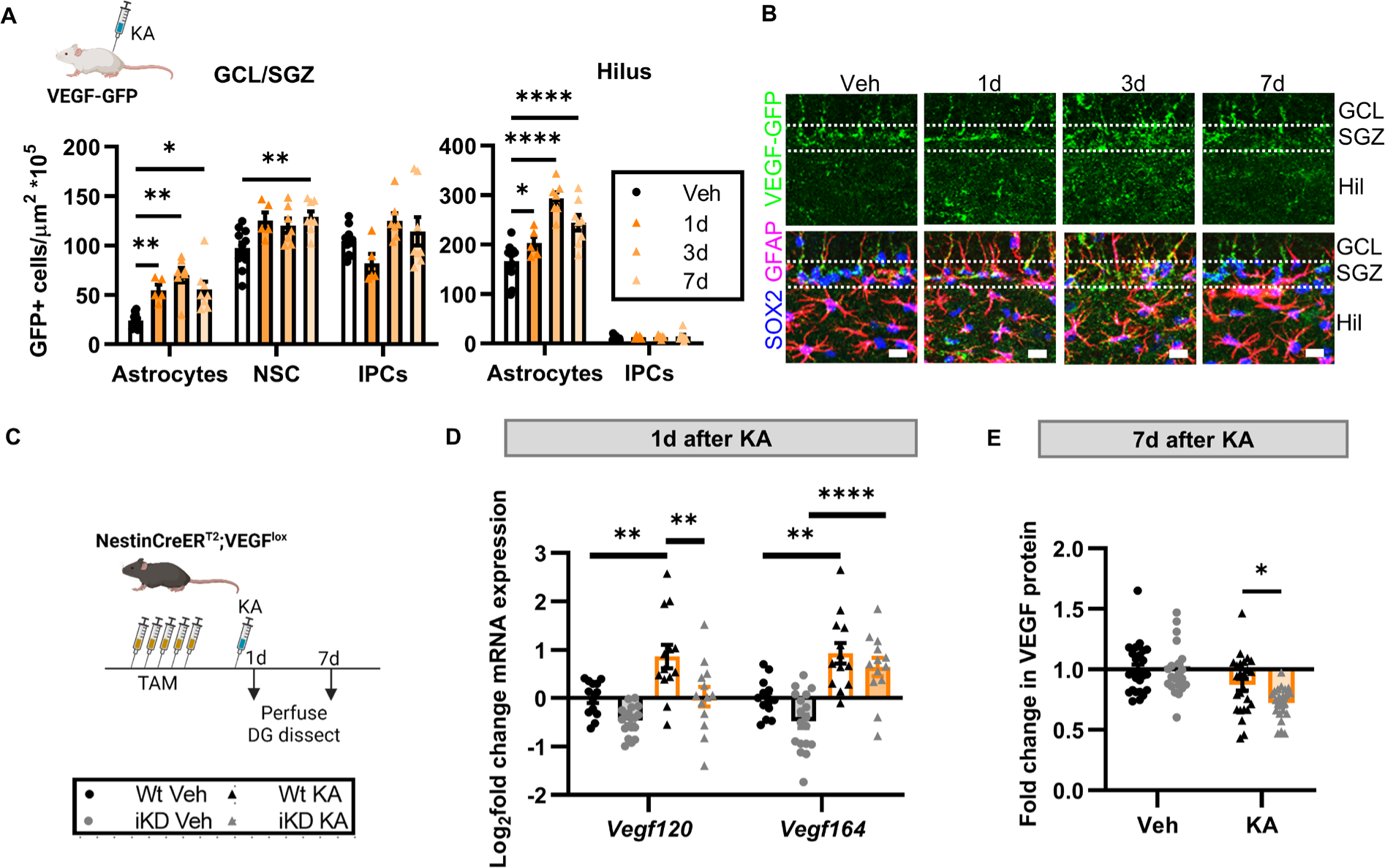
NSPCs contribute VEGF to the DG in response to excitotoxic injury. A) Density of VEGF-GFP+ cells in the granule cell layer/subgranular zone (GCL/SGZ) and hilus identified as astrocytes, NSCs or IPCs based on immunolabeling for Sox2 and GFAP quantified 1d, 3d, or 7d after a single systemic injection of KA in adult VEGF-GFP mice. 2 way repeated measures ANOVA: SGZ/GCL: cell type x timepoint p = 0.0053, cell type p < 0.0001, time p = 0.0002. Hilus: cell type x timepoint p < 0.0001, cell type p < 0.0001, time p < 0.0001. ******* p < 0.05, 0.01, 0.0001 Dunnett’s multiple comparisons to Vehicle. Mean ± SEM and individual mice shown. N = 11 vehicle, N = 5-8/KA timepoint. B) Representative images of VEGF-GFP, GFAP, Sox2 immunolabeling in the GCL, SGZ and hilus. Scale = 20 µm. C) Schematic of treatments. Wt and İKD mice received 5d TAM then 3-5 days later a single injection of systemic KA. DG was dissected 1 or 7d after KA. D) l_og_2_ fold change in Vegfa isoform mRNA relative to Wt-Veh control in the DG 1 d after KA. Three-way ANOVA isoform x treatment interaction p = 0.0231, treatment p < 0.0001, genotype p = 0.0008, isoform p = 0.0297. **,****p<0.01, 0.0001 Sidak’s multiple comparisons. Mean ± SEM and individual mice shown. N = 13-18 mice/group. E) Fold change in VEGF protein in whole DG relative to Wt-Veh control 7 d after KA. 2 way ANOVA treatment p <0.0001. *p = 0.0205, posthoc Sidak’s comparisons. Mean ± SEM and individual mice shown. N = 21-25 mice/group.

We next asked whether the proportion of total DG VEGF produced by NSPCs changes with KA. We measured VEGF mRNA and protein in whole DG 1 or 7d after KA in VEGF iKD and Wt mice (**Fig. 4C**). At 1d after KA, KA increased *vegf120* and *vegf164* mRNA by ∼2-fold in Wt mice. iKD mice showed significant blunting of the KA-response specifically in the *vegf120* isoform, but not in the *vegf164* isoform (**Fig. 4D**). At 7d after KA, DG VEGF protein (including both 120 and 164 isoforms) was significantly lower in iKD mice compared to Wt mice (**Fig. 4E**). Combined, these findings suggest that NSPCs contributed measurable amounts of VEGF to total DG VEGF, particularly of the *vegf120* isoform, both during an acute VEGF surge and a later recovery phase.

### Knockdown of NSPC-derived VEGF exacerbates excitotoxic injury

We next examined the injury response to KA-induced excitotoxicity. Using wildtype mice, we found that KA caused large increases in expression of proinflammatory genes *c1qa*, *il1a* and *tnf* that peaked between 1 and 3d after KA (**Supplementary Fig. 4A-D**). Expression of these genes, especially *tnf*, was most consistently high in the DG and CA regions of the hippocampus, with the cortex and subventricular zone (SVZ) showing no significant response to KA. This relative isolation of injury response to hippocampal regions is consistent with numerous observations of the hippocampus showing heightened sensitivity to injury relative to other brain regions^36^. We also confirmed that KA induced increases in DG microgliosis as revealed by immunolabeling for CD68 (**Supplementary Fig. 4E,F**) and astrogliosis as revealed by immunolabeling for GFAP (**Supplementary Fig. 4G,H**), both of which peaked 3-7d after KA.

Based on the above results, we first examined VEGF iKD and Wt mice 1d after KA to quantify the early phase of excitotoxic response (**Fig. 5A**). VEGF iKD and Wt mice both showed motor seizure activity after KA, with a mode Racine score of 3 for both groups (**Supplemental Fig. 5A**). cFos labeling in the granule cell layer, a surrogate measure of neuronal activity, was 66.6% greater in VEGF iKD mice than Wt littermates (**Fig. 5B,C**). Mice were also injected with BrdU during TAM injections, to mark dividing cells. Almost no BrdU+ cells co-labeled with cFos in any group, suggesting that cells born during TAM-induced recombination were not yet functionally integrated in circuitry, as would be expected based on their cellular age (**Supplemental Fig. 5B**). These findings suggested that loss of NSPC-derived VEGF causes an acute increase in excitability of existing neurons in the DG.

**Figure 5.**
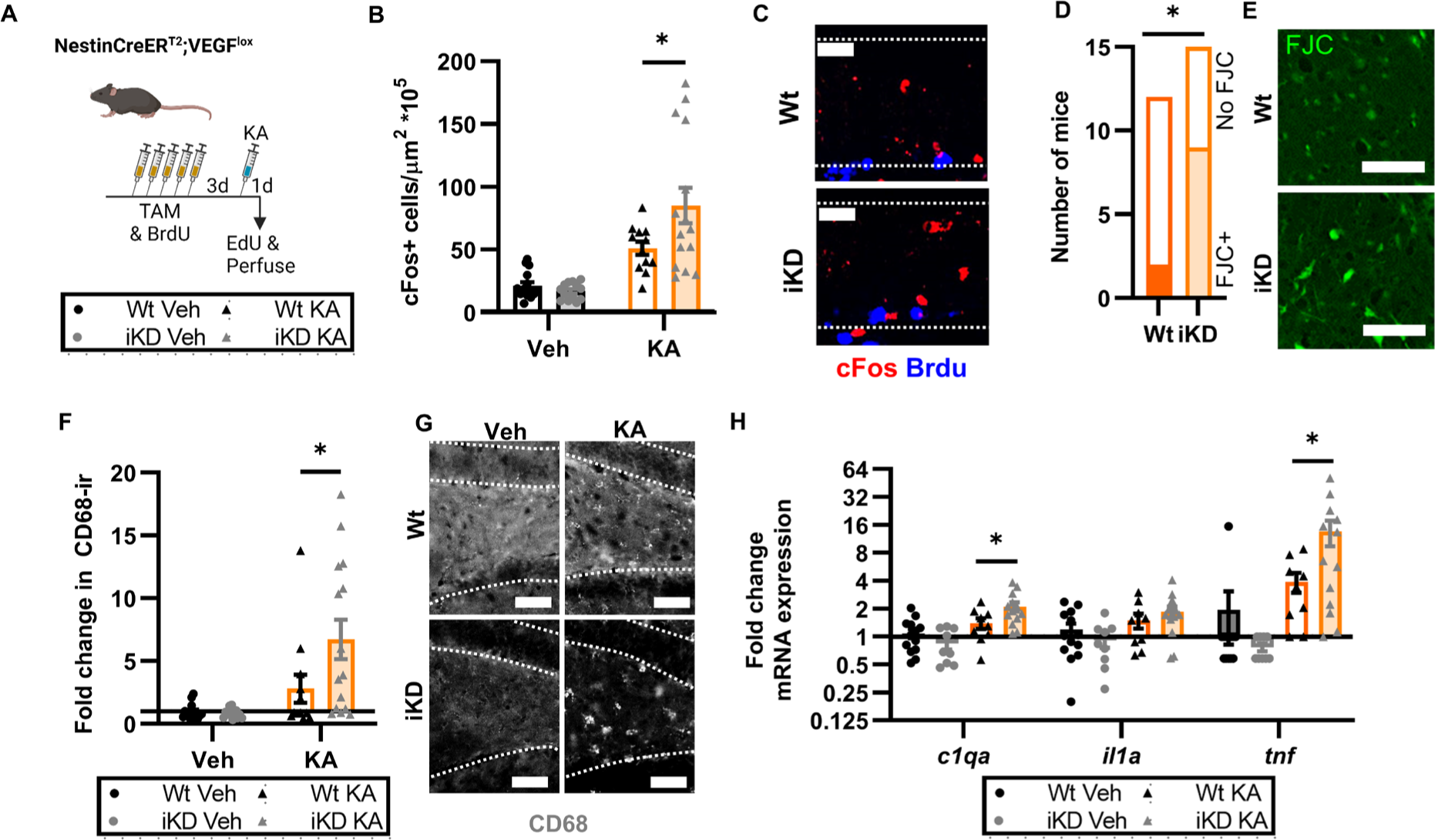
NSPC-VEGF knockdown exacerbates acute excitotoxic injury. A) Schematic of treatments. Wt and İKD mice received 5d TAM/BrdU then 3 days later a single injection of systemic KA. Mice were perfused 1d later. B) Density of DCX+ and DCX-cFos+ cells in the granule cell layer 1d after KA injection in Wt and İKD mice. 2 way ANOVA genotype x treatment interaction p = 0.0241, treatment p < 0.0001. *p = 0.0116 Sidak’s multiple comparisons. C) Representative images of immunolabeling for cFos and BrdU in the granule cell layer (dashed outline). Scale = 10 µm. D) Number of Wt and İKD mice showing any FJC labeling in the DG 1 d after KA. *p = 0.0473 Fisher’s exact test. E) Representative images of FJC labeling in the hilus (where almost all FJC+ cells were found) in one Wt and one İKD mice which showed FJC. Scale = 100 µm. F) Fold change in CD68 immunoreactivity (ir) thresholded area in the DG of Wt and İKD mice 1d after KA relative to Wt-Veh mice. 2 way ANOVA genotype x treatment interaction p = 0.0455, treatment p <0.0001. *p = 0.0161 Sidak’s multiple comparisons. G) Representative images of CD68ir in the DG of Wt and İKD mice. Granular cell layer has dashed outline. Scale = 20 µm. H) Fold change in mRNA expression of *c1qa, ill a* and *tnf* in whole DG in Wt and İKD mice 1d after KA injection relative to Wt-Veh mice. 2 way ANOVA: c1qa genotype x treatment interaction p = 0.0147, treatment p < 0.0001; ill a treatment p <0.0001; tnf genotype x treatment interaction p = 0.0468, treatment p = 0.0081. *p < 0.05 Sidak’s multiple comparisons. B,F,H) Mean ± SEM and individual mice shown. N = 12-15 mice/group.

We next used Fluorojade C labeling to quantify neuronal degeneration. Fluorojade C was only detected in DG of KA treated mice, with VEGF iKD mice being significantly more likely to show Fluorojade C+ degenerating cells than Wt littermates: 16.6% (2/12) of Wt mice treated with KA had detectable Fluorojade C in the DG while 60% (9/15) of VEGF iKD mice had detectable Fluorojade C (**Fig. 5D,E**). VEGF iKD mice also showed significantly greater signs of DG neuroinflammation in response to KA than Wt littermates as revealed by CD68 immunolabeling (**Fig. 5F,G**) and expression of *c1qa* and *tnf* (though not of *il1a*) (**Fig. 5H**). GFAP immunolabeling showed no change in response to KA at this 1 day timepoint (**Supplementary Fig. 5C,D**).

To confirm that KA did not alter the specificity of TAM-induced recombination, we quantified expression of stop-floxed EYFP in NestinCreER^T2^ mice 1d after KA. Almost all EYFP+ DG cells showed either NSC phenotype (56%) or IPC phenotype (41%), with <1% showing astrocytic phenotype (**Supplementary Fig. 5E-G**). In addition, to confirm that the apparent exacerbation of KA-induced injury due to loss of NSPC-VEGF was not an artefact of Cre expression, we injected VEGF^wt/wt^;NestinCreER^T2^ mice (Cre+) and VEGF^wt/wt^ littermates (Cre-) with TAM and KA. Cre+ and Cre-mice did not differ in Fluorojade C labeling or CD68 immunolabeling, suggesting that Cre expression alone does not drive exacerbation of excitotoxic injury in the DG (**Supplementary Fig. 5H,I**).

At 7d after KA, we found evidence of continued exacerbation of injury in VEGF iKD mice compared to Wt littermate controls (**Fig. 6A**). VEGF iKD mice showed greater frequency of Fluorojade C+ degenerating neurons (43% or 6/14) than in Wt mice (7% or 1/15) (**Fig. 6B,C**). CD68 immunoreactivity in the DG was no longer different between iKD and Wt mice (**Fig. 6D,E**), but GFAP immunoreactivity showed a KA-induced increase that was exacerbated in VEGF iKD mice compared to Wt mice (**Fig. 6F,G**). In agreement with previous work showing that excitotoxic injury causes vascular remodeling in the DG that can be reflected as reduced vessel coverage^37, 38^, we found that KA induced a loss of vessel coverage using immunolabeling for the endothelial cell marker CD31. However, this loss was similar in Wt and iKD mice (**Fig. 6H,I**), suggesting that gross vascular remodeling after KA injury is not dependent on VEGF derived from NSPCs.

**Figure 6.**
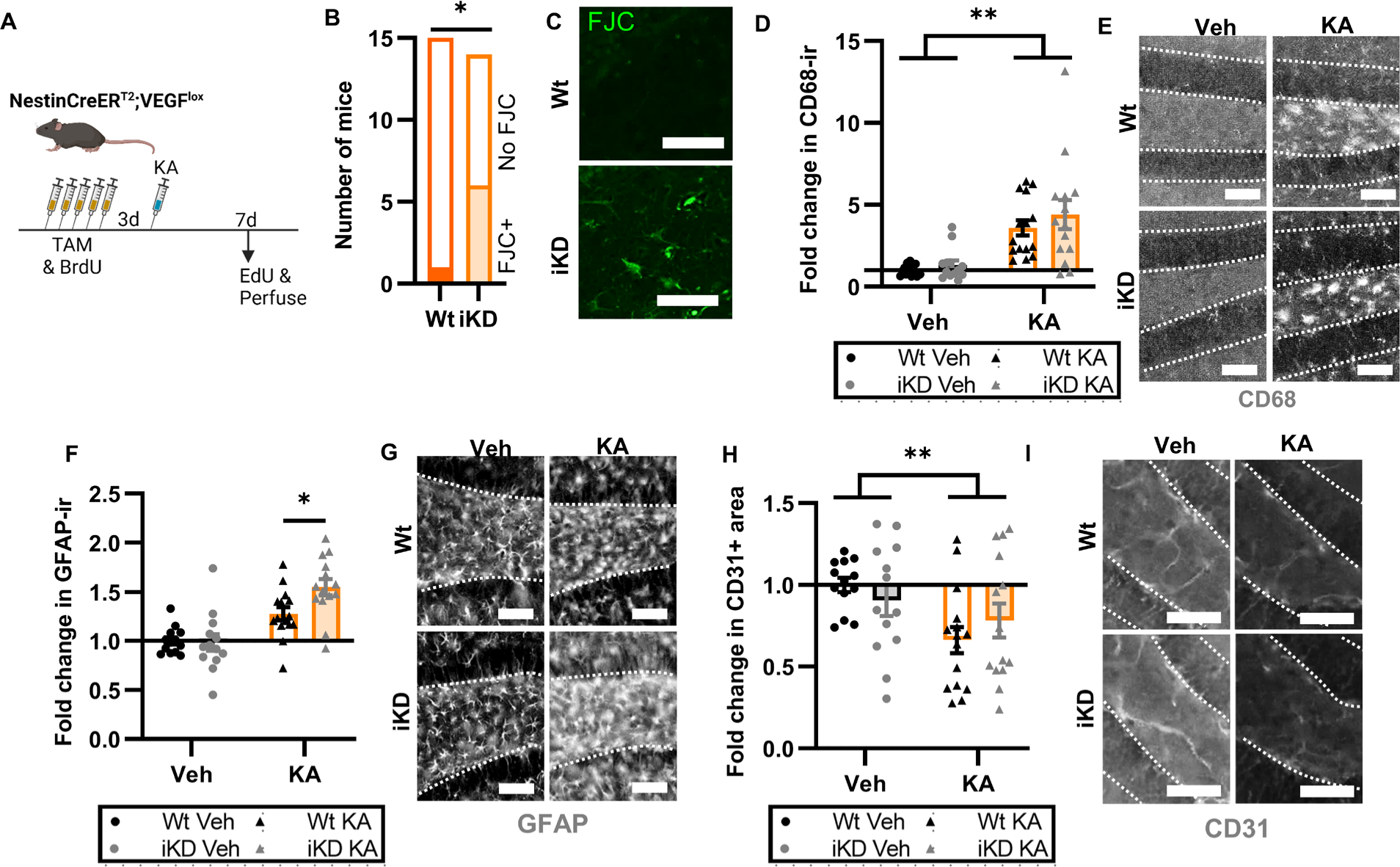
NSPC-VEGF knockdown exacerbation of excitotoxic injury persists 7d later. A) Schematic of treatments. Wt and İKD mice received 5d TAM/BrdU then 3 days later a single injection of systemic KA. Mice were perfused 7d later. B) Number of Wt and İKD mice showing any FJC labeling in the DG 7 d after KA. *p = 0.0352 Fisher’s exact test. C) Representative images of FJC labeling in the hilus (where almost all FJC+ cells were found). Scale = 100 µm. D) Fold change in CD68 immunoreactivity (ir) thresholded area in the DG of Wt and İKD mice 7d after KA relative to Wt-Veh mice. 2 way ANOVA “treatment p <0.0001. E) Representative images of CD68ir in the DG of Wt and İKD mice. Granular cell layer has dashed outline. Scale = 20 µm. F) Fold change in GFAP immunoreactivity (ir) thresholded area in the DG of Wt and İKD mice 7d after KA relative to Wt-Veh mice. 2 way ANOVA genotype x treatment interaction p = 0.0468, treatment p <0.0001. *p = 0.0151 Sidak’s multiple comparisons. G) Representative images of GFAPir in the DG of Wt and İKD mice. Granular cell layer has dashed outline. Scale = 20 µm. H) Fold change in CD31 immunoreactivity (ir) thresholded area in the DG of Wt and İKD mice 7d after KA relative to Wt-Veh mice. 2 way ANOVA treatment **p = 0.0094. I) Representative images of CD31ir in the DG of Wt and İKD mice. Granular cell layer has dashed outline. Scale = 20 µm. D,F,H) Mean ± SEM and individual mice shown. N = 13-15 mice/group.

Combined, we found substantial acute and long-lasting exacerbation of neuronal degeneration and neuroinflammation in the DG of VEGF iKD mice compared to Wt littermates in response to excitotoxic injury.

## Discussion

The stem cell secretome has recently emerged as a potent mechanism by which endogenous and transplanted stem cell populations can regulate adult tissue physiology. As efforts to harness stem cells as therapeutics advance, this functional dimension requires consideration^3, 8, 39^. Here, we demonstrate a beneficial role of endogenous hippocampal NSPC-derived VEGF in promoting hippocampal memory function and protecting against excitotoxic damage secondary to seizures, likely by suppressing neuronal excitability in DG circuitry.

Several other components of the DG NSPC secretome have been reported, primarily with functional roles in regulating NSPCs or neurogenesis^40^. For example, NSPCs in the adult hippocampus have been shown to secrete proteins that support either their own proliferative maintenance (e.g. IGF^41^, VEGF^13^, MFGE8^11^) or the maturation of immature neuronal populations (e.g. PTN^10^). Less is known about functional paracrine signaling from endogenous NSPCs. One recent study showed that NSPCs in the other major adult neurogenic niche, the SVZ, support striatal neuronal function via secretion of IGFBPL1^12^. We have previously shown that NSPCs in the SVZ do not synthesize VEGF^13^, making it unlikely that the cells in this other niche use VEGF similarly to what we found in the DG. However, these complimentary findings with IGFBPL1 further highlight the potential for direct modulation of local neuronal function by an endogenous NSPC secretome. More research is needed to better identify other components of the endogenous NSPC secretome and functional relevance.

Previous research presents often opposing findings about how VEGF affects hippocampal function. These opposing findings have garnered VEGF a reputation as a “double-edged sword” ^42^, particularly in the context of excitotoxic injury. Generally, in response to hippocampal injury, VEGF signaling to neurons is considered neuroprotective, while its effects on vascular cells is considered detrimental^14^. Our findings suggest that VEGF derived from endogenous NSPCs is beneficial to hippocampal memory function and suppresses hyperexcitation. This beneficial effect extends to the response to excitotoxic injury, providing protection against neuronal injury and gliosis. We did not detect any effect of NSPC-derived VEGF on vascular remodeling after excitotoxic injury, suggesting that VEGF from another source (or other factors) mediate this vascular response. Future research will be necessary to determine the mechanisms underlying DG vascular remodeling after excitotoxicity.

Historically, VEGF production in the brain has been attributed to astrocytes^43^. Here, we show that, while astrocytes are the majority VEGF-expressing population by cell number in the DG, NSPCs still contribute substantial quantities of VEGF, with the most notable contribution being to the *vegf120* isoform in response to kainic acid excitotoxicity. This finding is consistent with our previous finding that NSPCs synthesize both *vegf120* and *vegf164*^13^, while astrocytes are known to synthesize *vegf164* primarily^44^. We also found that both major VEGF isoforms (VEGF_120_ and VEGF_164_) had wide and generally similar dispersal properties in mouse DG in vivo. The wide dispersal of VEGF protein that we observed suggests that the restriction of NSPCs to the subgranular zone is not a likely impediment to NSPCs affecting DG cells broadly via secreted VEGF. It also further suggests that VEGF derived from stem cell transplants may have the potential to affect large portions of hippocampal tissue, even if cells do not migrate long distances themselves.

VEGF_120_ and VEGF_164_ differ primarily in their relative solubility. VEGF_164_ carries C-terminal heparin-binding domains that partially bind in the extracellular matrix, while VEGF_120_ does not^45^. As a result, VEGF_120_ is frequently described as soluble while VEGF_164_ is described as semi-soluble. The similarity in dispersal between VEGF isoforms that we observed here therefore seems in opposition to these known solubility differences. However, computational estimates reveal that the dominant limiter of VEGF dispersal in vivo is binding and sequestering by VEGF receptors, not isoform solubility^28^. These two VEGF isoforms have similar receptor binding capabilities, possibly explaining the general similarities in their dispersal and clearance characteristics in the present study^44^.

Our findings of neuroprotection via NSPC-VEGF add a new dimension to consider when investigating the role of adult neurogenesis in seizure response. Parent et al.^46^ originally described aberrant neurogenesis in the adult rodent DG after induced seizures. They reported a surge in production of new neurons that migrated to the hilus, rather than the granule cell layer, and sent axonal projection to inappropriate DG cell layers. These findings led to the hypothesis that aberrant newborn neurons may contribute to development of epileptic circuits in the hippocampus, a hypothesis that has recently been supported by findings in humans^47^.

Nonetheless, contrasting rodent studies suggest a potentially complex role for new neurons in seizures, with some neuronal subsets contributing to epileptogenesis and others preventing it^48, 49^. Our findings add this growing literature, suggesting that NSPCs themselves play a beneficial role in protecting the DG. Attempts to harness endogenous neurogenic processes in any kind of therapeutic intervention for epilepsy will benefit from considering these multiple functional domains of the NSPCs and their progeny.

There are several limitations of the present study. First, though we identify neuronal hyperexcitability after NSPC-VEGF loss, we do not identify whether this happens via direct VEGFR2 signaling on excitatory granule cells, via direct VEGFR signaling on another neuronal subtype, or via signaling to another non-neuronal intermediary cell. Knockdown of VEGFR2 in hippocampal neurons impairs memory function^22, 23^, making a direct loss of VEGFR2 signal a likely mechanism, but more research is needed to confirm this hypothesis. A challenge moving forward will be how to distinguish effects from NSPC-derived VEGF versus VEGF derived from other cells (e.g. astrocytes) on target cells/receptors. Second, our estimates of VEGF dispersal in vivo derive from an artificial source of VEGF and therefore may differ from how VEGF disperses when secreted from endogenous cells. A more accurate assessment would require tagging of proteins synthesized in vivo, which remains a challenge for the future.

In summary, we show that NSPCs support and protect the hippocampus from injury via production of VEGF. These findings provide a functional role for NSPCs independent of their capacity to generate new neurons. This new functional role is worthy of consideration in attempts to develop therapies leveraging endogenous NSPCs. It also warrants consideration within the ongoing debate about the existence of adult neurogenesis in humans. Much of this debate has focused on generation of new neurons as the crucial functional output of the neurogenic lineage^7, 50–54^. The present work adds to an emerging literature suggesting that NSPCs themselves may have functional roles which could persist even in the absence of generation of new neuronal progeny.

## Methods

### Animals

All mice were group housed (2-5 same sex mice/cage) with food and water ad libitum with a 12h light-dark cycle, lights on at 0630h. Males and females were used in all experiments in approximately equal numbers. VEGF^lox^ mice were provided by Genentech Inc^17^. NestinCreER^T2^ mice, previously described^16^, were obtained from Jackson Labs (strain #01621). VEGFGFP mice were a gift from Brian Seed, Harvard University, Cambridge, MA^55^. Rosa-Stopflox-EYFP mice were obtained from Jackson labs (strain #006148). Transgenic mice were all bred in-house and used for experiments at 5-9 weeks of age. Wt C57Bl6/J mice were obtained from Jackson Labs, (strain #000664) at 6 weeks of age and used for experiments at 7-8 weeks of age. Genotyping primers were: VEGF^lox^(by gel electrophoresis) F 5′cctggccctcaagtacacctt, R 5′- tccgtacgacgcatttctag; Cre (by gel electrophoresis) F 5′-gcggtctggcagtaaaaactatc, R 5′- gtgaaacagcattgctgtcactt; Cre (by real time qPCR) F 5’-cggtctggcagtaaaaactat, R 5’- cagggtgttataagcaatccc; GFP (by gel electrophoresis or real time qPCR) F 5′- tccttgaagaagatggtgcg R 5′-aagttcatctgcaccaccg; EYFP (by gel electrophoresis) F 5′- aaagtcgctctgagttgttat, R (Wt) 5′-ggagcgggagaaatggatatg, R (mut) 5′-aagaccgcgaagagtttgtc.

### Animal treatments

TAM (Fisher, # 50-115-2413) was dissolved (20 mg/ml) in sterile sunflower oil (Fisher, # 18-607-188) overnight at 37°C with rotation in the dark then stored at +4°C for 1 week maximum. TAM was injected intraperitoneally at 180 mg/kg for 5d. Kainic acid (Sigma, # 420318) was dissolved (5 mg/ml) in sterile saline and stored at −20°C. We used 15-20 mg/kg KA delivered intraperitoneally to stimulate overexcitation without triggering status epilepticus (which has high fatality in C57Bl6 mice). BrdU (Sigma, # B5002) and EdU (Fisher, # NC1296287) were dissolved in sterile saline (10 mg/ml) and injected intraperitoneally 150 mg/kg/d.

### Perfusion and tissue harvest

For euthanasia, mice were anesthetized with ketamine (87.5 mg/kg) and xylazine (12.5 mg/kg) followed by transcardial perfusion with 0.01M phosphate buffered saline (PBS).

### Immunolabeling

Immunolabeling was performed similar to our previous work^18, 56^. After perfusion, brains were harvested and fixed 24h in 4% paraformaldehyde in 0.1 M phosphate buffer followed by equilibration in 30% sucrose in PBS, both at 4°C. Brains were sliced on a freezing microtome (Leica) in a 1 in 12 series of 40 µm slices and stored in cryoprotectant at −20°C. Slices were rinsed, incubated in blocking solution (1% normal donkey serum, 0.3% triton X 100 in PBS) for 30 min at room temperature then transferred to primary antibody in blocking solution overnight at +4°C with rotation. The next day, slices were washed and incubated in secondary antibody (1:500) in blocking solution for 2h at room temperature with rotation, followed by 10 min in Hoechst (1:2000 in PBS, Fisher #H3571). After rinsing, slices were mounted on superfrost plus microscope slides precleaned (Fisher, #12-550-15) and protected with Prolong Gold antifading medium (Fisher, #P36934) with coverglass. For BrdU labeling, all other antigens were labeled first, followed by 10 min in 4% paraformaldehyde, rinsing and 30 min in 2N HCl at 37°C. Sections were then rinsed blocked and labeled for BrdU. For EdU labeling, click labeling was performed according to manufacturer instructions (ClickChemistryTools) followed by immunolabeling. Antibodies are summarized in the Table below.

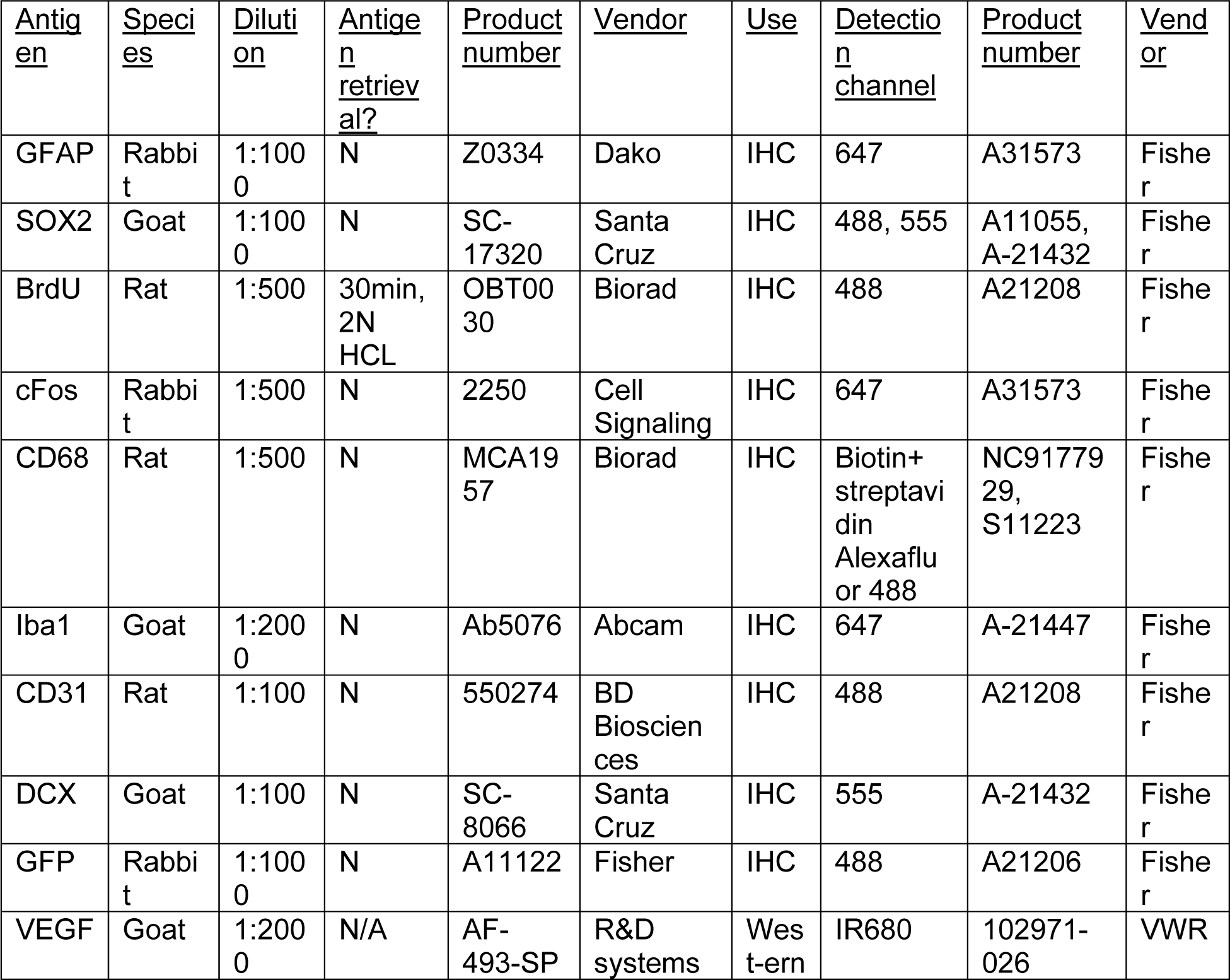

| <u>Antigen</u> | <u>Species</u> | <u>Dilution</u> | <u>Antigen retrieval?</u> | <u>Product number</u> | <u>Vendor</u> | <u>Use</u> | <u>Detection channel</u> | <u>Product number</u> | <u>Vendor</u> |
| --- | --- | --- | --- | --- | --- | --- | --- | --- | --- |
| GFAP | Rabbit | 1:1000 | N | Z0334 | Dako | IHC | 647 | A31573 | Fisher |
| SOX2 | Goat | 1:1000 | N | SC-17320 | Santa Cruz | IHC | 488, 555 | A11055, A-21432 | Fisher |
| BrdU | Rat | 1:500 | 30min, 2N HCL | OBT0030 | Biorad | IHC | 488 | A21208 | Fisher |
| cFos | Rabbit | 1:500 | N | 2250 | Cell Signaling | IHC | 647 | A31573 | Fisher |
| CD68 | Rat | 1:500 | N | MCA1957 | Biorad | IHC | Biotin+ streptavidin Alexafluor 488 | NC9177929, S11223 | Fisher |
| Iba1 | Goat | 1:2000 | N | Ab5076 | Abcam | IHC | 647 | A-21447 | Fisher |
| CD31 | Rat | 1:100 | N | 550274 | BD Biosciences | IHC | 488 | A21208 | Fisher |
| DCX | Goat | 1:100 | N | SC-8066 | Santa Cruz | IHC | 555 | A-21432 | Fisher |
| GFP | Rabbit | 1:1000 | N | A11122 | Fisher | IHC | 488 | A21206 | Fisher |
| VEGF | Goat | 1:2000 | N/A | AF-493-SP | R&D systems | Western | IR680 | 102971-026 | VWR |

### Microscopy and image quantification

All images using co-labeling were acquired on a Zeiss Axio Observer Z1 microscope with Apotome for optical sectioning or a confocal laser-scanning microscope (Zeiss) using 20x air, 40x air or 63x oil objective as needed. Full z-stacks were acquired for analysis. Quantification of astrocytes, NSCs and IPCs was done by 1-2 blinded observers in z-stacks to identify GFAP+SOX2+ stellate cells (astrocytes), GFAP+SOX2+ radial glia like cells (NSCs) and SOX2+ cells lacking GFAP co-label (IPCs). Cell counts were corrected for area sampled as appropriate (SGZ, GCL, Hilus as described in figures) to yield a density per area. Images of Fluorojade C, CD68, Iba1, GFAP and CD31 as single labels were acquired using an automated slide scanner (Hamamatsu Photonics). CD68, Iba1, GFAP and CD31 single labels were all quantified using thesholded area analysis in ImageJ. For RNAscope analysis, Hoechst 33342/GFAP were used to identify stellate astrocytes and radial glia-like NSCs then Vegfa puncta were counted inside the nucleus of each cell.

### Generating tagged VEGF constructs

VEGF_120_ and VEGF_164_ were synthesized by Genscript using the known sequences of mouse VEGF_120_ and VEGF_164_ cDNA. A TAG stop codon inserted at bp100-102, corresponding to the location after the 33^rd^ amino acid, within the second exon. VEGF120 and VEGF164 have identical sequences through this point in their coding sequence. This location was chosen because this is the location of an additional amino acid in human VEGF compared to mouse VEGF isoforms. Human and mouse isoforms of VEGF show great functional overlap, leading us to hypothesize that an insertion here would not disrupt the conformation of mouse VEGF. Human VEGF protein sequence from signal peptide (in parentheses) to amino acid 40 is (MNFLLSWVHW) SLALLLYLHH AKWSQAAPMA EG**G<u>G</u>Q**NHHEV. The parallel mouse VEGF protein sequence is (MNFLLSWVHW) TLALLLYLHH AKWSQAAPTT EG**EQ**KSHEVI. The TAG-inserted constructs would code for (MNFLLSWVHW) TLALLLYLHH AKWSQAAPTT EG**E*Q**KSHEVI, where * indicates the TAG amber stop codon. These constructs were inserted downstream of an Ef1α promoter in a viral expression backbone, with a 6x His tag added at the end of the coding sequence. These constructs are in the process of being deposited at Addgene for public use.

### Synthesis of VEGF^TAG^ proteins

Human embryonic kidney (HEK293T) cells (ATCC, #CRL-3216) were cultured in standard conditions on uncoated flasks: DMEM with high glucose, pyruvate, GlutaMAX (Fisher, #10-569-010) with 10% FBS (Fisher, #10-437-028), 1x MEM non-essential amino acids (Fisher, #11-140-050) and Pen/Strep (Fisher, #15140-122). In commercial production, recombinant proteins are often synthesized using bacterial cells. However, post-translational modifications can differ greatly between prokaryotes and eukaryotes. We therefore chose to use mammalian cells (HEK cells) to generate proteins that were as similar to endogenous mammalian VEGF isoforms as possible. Cells were transfected using lipofectamine 3000 (Fisher, #L3000015) according to manufacturer instructions with: VEGF_120_^TAG^, VEGF_164_^TAG^ and/or PylRS/tRNApyl (Addgene #105830 from^29^). TCO*A (SiChem, #SC-8008) was diluted 1:4 in 10X PBS then spiked into culture dishes to yield a final concentration of 1 mM. One day later, media was changed to serum-free media with 1 mM TCO*A. 1 day later, CM was collected and concentrated 10x with Amicon Ultra 3 kDa filter (Millipore, #UFC900308) according to manufacturer instructions. We chose to isolate protein from CM to capture the secreted form of VEGF after all post-translational modifications prior to secretion were complete. CM concentrate was incubated with High-Capacity Ni-IMAC Magnetic Beads (Invitrogen, # A50588) to select His-tagged proteins according to manufacturer instructions. Bead supernatant was click reacted with tetrazine-biotin (50 µM) then dialyzed 3h and then overnight in 0.01 M PBS using G2 Dialysis Cassettes 3.5K MWCO (PI87722, Fisher) to eliminate free tetrazine-biotin. Dialysate was then concentrated with Amicon Ultra 3 kDa filter (Millipore, #UFC900308) according to manufacturer instructions and stored at −80°C until use. Vehicle control was CM from HEK cells transfected with PylRS/tRNApyl only and processed in parallel with tagged CM samples.

To confirm the efficacy and specificity of tagging, samples were separated by PAGE in 4–15% Mini-PROTEAN® TGX™ Precast Protein Gels (BioRad, #4561084) for 1hr at 120V in Tris/Glycine/SDS running buffer (BioRad, #161-0772). Proteins were transferred to nitrocellulose overnight in 20% methanol in Tris/Glycine transfer buffer (VWR, #97061-382). Total protein was visualized using the LI-COR REVERT™ Total Protein Stain (Licor, #926-11021) with IR imaging on a Licor Odyssey Clx (Licor). Total protein was reversed, and membranes were reacted with streptavidin-IR800 (VWR, #103011-496) for 1h at room temperature, followed by rinsing in TBS-t and blocking in 5% milk in TBS-t for 1h at room temperature. Membranes were then incubated in primary antibody in blocking solution overnight at 4°C. The next day, membranes were rinsed 3x in TBS-t, incubated in secondary antibody diluted 1:10,000 in blocking solution and rinsed 3 more times before visualization on a Licor Odyssey Clx imager.

### Stereotaxic infusion of VEGF^TAG^ proteins

Adult male and female C57Bl6 mice were anesthetized by isoflurane inhalation (Akorn, 4% induction, 2% maintenance) in oxygen. After mounting in a stereotaxic frame (Stoelting), alcohol and betadine were used to sterilize the scalp. After exposing the skull using a scalpel blade, a unilateral hole was drilled at approximate 2.0 mm posterior to bregma. Hamilton syringes were lowered to the following position: A/P −2.0 mm, M/L +2.0 mm, D/V −1.7 mm relative to bregma. 2.5 ng of VEGF^TAG^ or matched volume of vehicle control sample was infused unilaterally (right and left side counterbalanced across animals) at a rate of 0.1 µl/min using an automated injector (Stoelting). Mice were administered carprofen prior to surgery. If mice were in the 1h, 3h or 6h groups, they also received buprenorphine immediately post-surgery. At their designated timepoint, mice were perfused as described above. A total of 59 animals received unilateral infusion (Veh: 12 mice; VEGF_120_^TAG^: 24 mice, and VEGF_164_^TAG:^ 23 mice).

### Detection and quantification of VEGF^TAG^ proteins in vivo

Fixed brains were sliced into 40 µm coronal slices and stored in cryoprotectant at −20°C. Slices were later rinsed 3x with PBS before mounting the entire rostral-caudal extent of the hippocampus on superfrost plus microscope slides precleaned (Fisher, #12-550-15). After drying overnight, slices were incubated in streptavidin-IR800 (VWR, #103011-496) diluted 1:2000 in PBS for 1 hr then rinsed 3 more times in PBS before drying and coverslippping with Invitrogen ProLong Gold Antifade Reagent (Fisher, #P36934). Slides were imaged on a Licor Odyssey Clx imager. Area of IR800 signal was detected and measured using the Licor Small Animal Imaging Analysis key (Licor, #2000-502), with detection thresholds set at standard deviation 4.0 relative to tissue background and search limit 50 pixels. The needle track was used to set the epicenter of the infusion, which included the observed center of the track plus 4 slices rostral and caudal. To correct for area detected from tissue autofluorescence in the DG, vehicle sample average area per slice was subtracted from all areas to express signal beyond background. Of the 59 animals infused, 56 survived to perfusion and needle tracks were confirmed to be in the hippocampus for 42 mice: Veh: 2 10 min mice, 2 1h mice, 2 3h mice, 3 6h mice; VEGF_120_^TAG^: 4 10 min mice, 4 1h mice, 4 3h mice, 4 6h mice, and VEGF_164_^TAG:^ 5 10 min mice, 4 1h mice, 4 3h mice, 4 6h mice. Vehicle mice showed no differences across timepoints and were collapsed into a single group for analysis.

### RNAscope© in situ hybridization

3d after the last TAM injections, Wt and VEGF iKD mice were transcardially perfused with ice cold PBS and 4% paraformaldehyde. Tissue processing is further described in^57^. Briefly, brains were harvested and fixed overnight in 4% paraformaldehyde followed by serial equilibration in 10-30% sucrose. Tissue was snap frozen in OCT and stored at −70°C until sectioning in to 12 µm slices on a cryostat (Leica). Thaw-mounted sections were stored at −70°C. RNAscope Multiplex Fluorescent v2 Assay (Advanced Cell Diagnostics, Newark, CA, United States) was performed with probe against mouse Vegfa (Mm-Vegfa-ver2, ACD 41226) according to manufacturer recommendations followed by immunolabeling for GFAP. Immunolabeling was similar to that described above with the following exceptions: blocking = 10% normal donkey serum in TBS-1% BSA, antibody diluent = TBS-1% BSA, washes = TBS-t.

### Behavioral testing

Mice were all handled by experimenters for at least 3 days before beginning testing. Both male and female experimenters were used and the same team members habituated the mice to handling and ran behavioral tests. The Y-maze test was conducted using an opaque, light-grey, arena made of extruded acrylic sheets with three identical arms of equal dimension (37 cm × 6 cm × 13 cm) angled 120° away from each other from a central point. Outside the arena, spatial cues were set up on each of the four sides of the rectangular table upon which the maze was set up. Mice were released into a designated “release arm,” and an opaque curtain was set up between the testing area and the experimenters. Four mice were trained or tested at once, with a camera set up centrally on the ceiling. During the Y-maze training (15 min), mice were able to roam freely in the release arm and one other arm, with access to the third arm blocked by a rectangular insert made of the same material as the arena. During the ITI (2 h), mice were returned to their experimental cages and placed back into the procedure room. Following the ITI, mice were returned to the Y-maze for 5 min with access to all three arms open. Once the test was complete, mice were once again returned to their home cages. Between each group of mice, each Y-maze arena was thoroughly wiped with 70% alcohol and completely dried.

After Y maze testing, mice were habituated to an open field, which was an opaque, light-grey, rectangular arena (40 cm × 39 cm × 30 cm) made of extruded acrylic sheet. During the habituations (3 x 5 min), each mouse was released into the designated release corner of the arena, with access to the entire, empty space. Four mice were habituated at once, with a camera set up centrally on the ceiling. Between each habituation, each spent a 50-minute ITI singly housed in their experimental cage in the procedure room. Between each group of mice, each OLT arena was thoroughly wiped with 70% alcohol and completely dried. At the end of the third habituation, mice were returned to their original cages.

The day after Y maze and open field habituation, mice were returned to the open field for novel object location testing (OLT), which was performed similarly to our previous work^58^. During training, the arena contained two objects, each one 9 cm away from each of the two closest walls of the arena. A 6.5 cm-tall, plastic cone filled with orange sand and a 9 cm-tall, plastic elephant toy were used as the objects. Each mouse was released into a designated release corner into the same arena in which they were habituated and allowed to explore for 10 min. Four mice were trained or tested at once, with a camera set up centrally on the ceiling. During the ITI (1 h), mice were returned to their home cages and placed back into the procedure room. The arenas were cleaned and one object was moved to a new corner-adjacent location. After the ITI, mice were returned to the arenas and allowed to explore for 10 min. Mice were then returned to their home cages. Between each group of mice, each OLT arena and all objects were thoroughly wiped with 70% alcohol and completely dried.

After the OLT task, mice were placed in the closed arm of an elevated plus maze and allowed to explore for 10 min. The elevated plus maze consisted of four arms—two open (6 cm × 34.5 cm) and two enclosed (6 cm × 34.5 cm × 21.5 cm). The arms were angled 90° from a central platform (6 x 6 cm).

### Behavioral analysis

Behavior was scored manually by 1-2 blinded observers in video recordings of all tasks. In the Y maze, the duration of time spent in each arm, frequency of visits to each arm, and latency to investigate the novel arm were all scored manually. Percent time in the novel arm was calculated relative to total time novel and familiar arm (excluding time in the release arm). In OLT, investigation time with each object was scored manually. Object investigation was defined as a mouse’s nose being towards and within 2 cm of the object, as described in our previous work^58^. Climbing the object was not considered as object investigation. Percent investigation time was calculated as (investigation time of moved object) / (investigation time of both stationary and moved objects). In EPM, time in open arms and entries in to closed and open arms were scored manually. An entry into an arm was counted when the mouse placed two or more paws into the new area.

### Extracellular electrophysiology

Mice were anesthetized with isoflurane before decapitation. The brain was rapidly removed and placed in an ice-cold cutting solution containing the following (in mM): Sucrose 250, D-Glucose 25, KCl 2.5, NaHCO_3_ 24, NaH_2_PO_4_ 1.25, CaCl_2_ 2.0, MgSO_4_ 1.5, and Kynurenic acid 1.0 (pH; 7.3-7.4). Transverse hippocampal slices (400 µm) were prepared with a Vibratome (VT1200S, Leica). Brain slices were transferred to chambers filled with artificial CSF (aCSF; bubbled with 95% O_2_ and 5% CO_2_) containing the following (in mM): NaCl 124, KCl 3, NaHCO_3_ 24, NaH_2_PO_4_ 1.25, CaCl_2_ 2, MgSO_4_ 1.0, and D-Glucose 10, (pH; 7.3-7.4). Slices were allowed to recover at 37°C for 30 min and then moved to room temperature for at least 1 hour before recording. Individual slices were transferred to a submerged chamber on the fixed stage of Nikon E600FN upright microscope. The chamber was perfused with oxygenated aCSF (containing 50 µM Picrotoxin) at 2-3 ml/min by gravity and maintained at 32°C during the recording.

Local field excitatory postsynaptic potentials (fEPSPs) were recorded from the molecular layer of the dentate gyrus with borosilicate glass electrodes (1.5-3 MΩ, filled with aCSF) and evoked by electrical stimulation of the afferent fibers of the medial perforant path (MPP) located in the upper blade of the DG. Stimulation pulses (100 µS duration, every 30 s) were generated by an isolator (Iso-flex, A.M.P.I.) under computer control and delivered through a custom-made twisted nichrome wire stimulating electrode. Input–output (I/O) curves were generated for each slice with stimulation intensity varying from 0-1.0 mA at each 0.1 mA step. The stimulation intensity which evoked ∼ 50% of the maximum response, without emergence of population spikes, was chosen to investigate LTP and the Paired-Pulse Ratio (PPR). The PPR were recorded at 6 time intervals (30, 50, 100, 150, 200, 250 mSec). Synaptic field potentials were low-pass filtered at 1 kHz and digitally sampled at 50 kHz with Axopatch 200B amplifier and Digidata 1440 A interface. The baseline was monitored for 20 min to establish a consistent response. High frequency stimulation consisted of four sets of 100 pulses, each delivered in 1 second (100Hz) and administered in 20 second intervals. The LTP recording lasted 60 minutes after high frequency stimulation. The slope (20-80%) of the evoked fEPSPs was measured and normalized to the average baseline 5 min prior to delivery of high frequency stimulation. Data were monitored on-line and analyzed off-line using Clampex 10.6 software.

### VEGF ELISA

After PBS perfusion, whole DG was dissected and flash frozen on dry ice then stored at −20°C. Protein was extracted from whole DG by homogenization in RIPA buffer (Fisher, #PI89900) with Halt Protease and Phosphatase Inhibitor (Fisher, #PI78440). Samples were incubated on ice for 10 min then freeze-thawed using dry ice and wet ice respectively 3 times. Samples were centrifuged at 14,000 rpm for 5 min at 4°C. Total protein content of supernatant was determined using a BCA kit (Fisher, #PI23225). 40 µg total protein was diluted in assay buffer and quantified using the R&D Systems Duo set VEGF ELISA according to manufacturer instructions.

### Fluorojade C labeling

Tissue was collected, sliced and stored as for Immunolabeling. Slices were rinsed then mounted on superfrost plus microscope slides precleaned (Fisher, #12-550-15). After drying overnight, slides were: immersed in 1% NaOH, 80% ethanol 5 min, followed by 70% ethanol for 2 min, 2×2min ddH2O rinses and 15min incubation in 0.06% KMnO4. After 2×2min ddH2O rinses, slides were immersed in 0.0001% Fluorojade C (Histochem, # 1FJC) in 0.1% acetic acid in the dark. After 3×2min ddH2O rinses, slides were allowed to dry overnight. After 2×2min rinses in xylenes, slides were protected with DPX mounting medium (Fisher, #50-980-369) and coverglass. Presence or absence of Fluorojade C in the DG was assessed by an observer blind to all treatments/genotypes.

### Real time qPCR

After PBS perfusion, whole DG was dissected and flash frozen on dry ice then stored at −80°C. RNA was extracted using the RNeasy Mini Kit (Qiagen, #74104) according to manufacturer’s instructions. RNA was DNase I (Invitrogen, # 18068015) treated and then converted to cDNA using the SuperScript III Frist Strand Synthesis System (Fisher, # 18080051). cDNA diluted 1:5 in water was quantified using either SYBR Green I (Roche, # 04707516001) and a LightCycler 480 (Roche) or SsoAdvanced-Universal SYBR Green Supermix (Biorad, # 1725274) and a BioRad CFX96 Touch Real-Time PCR Detection System. ΔΔCt values were calculated relative to housekeeping genes (actin and or sdha) and converted to log_2_fold change over control.

### Primers

C1qa F 5’- aaaggcaatccaggcaatatca, R 5’-tggttctggtatggactctcc (Primer bank 6671650a1) Il1a F 5’-gcaccttacacctaccagagt, R 5’-aaacttctgcctgacgagctt (Primer bank 6754328a1) Tnf F 5’- cctgtagcccacgtcgtag, R 5’-gggagtagacaaggtacaaccc (Primer bank 133892368c3) Vegf120 F 5’- agccagaaaaatgtgacaagc, R 5’- tctttccggtgagaggtctg from^13^ Vefg164 F 5’- agccagaaaatcactgtgagc, R 5’- gcgagtctgtgtttttgcag from^13^ Sdha F 5’- ggaacactccaaaaacagacct, R 5’-ccaccactgggtattgagtagaa (Primer bank 15030102a1) Actin F 5′-ggctgtattcccctccatcg, R 5′-ccagttggtaacaatgccatgt (Primer bank 6671509a1)

### Racine scoring

Mice were videotaped for 4h after KA injection and seizures were scored by a blinded observer according to the Racine scale: 1) mouth and facial movement, 2) head nodding, 3) forelimb convulsion, 4) Rearing plus forelimb clonus, 5) generalized motor seizure with rearing and falling.

### Statistical analysis

Statistical analysis was performed using GraphPad Prism. Specific tests used are detailed in figure legends. Sample sizes were always individual mice unless otherwise noted and are detailed in figure legends. In general, if 2 groups were compared, student’s t-tests or Mann-Whitney tests were used. If more than 2 groups were being compared within one factor, ANOVA was used with posthoc error corrected tests. If more than 2 groups were being compared with 2 or more factors, 2 way ANOVA was used with posthoc error corrected tests. Y maze arm first arm entry was compared using Fisher’s exact test. Spatial dispersal of tagged VEGF was compared using 95% confidence intervals because we had no apriori hypothesis in this experiment. P<0.05 was defined as significant.

## Acknowledgements

The authors would like to thank Tony Wyss-Coray for feedback and support in initiating this work. This work was funded by R00NS089938 and R01NS124775 to EDK. MH was funded by the CIRM BRIDGES program. TD and RR were each funded by Ohio State Undergraduate Research Scholarships. FZ and CA were funded by P30NS104177.

## Supplemental Figures

**Supplemental Figure 1.**
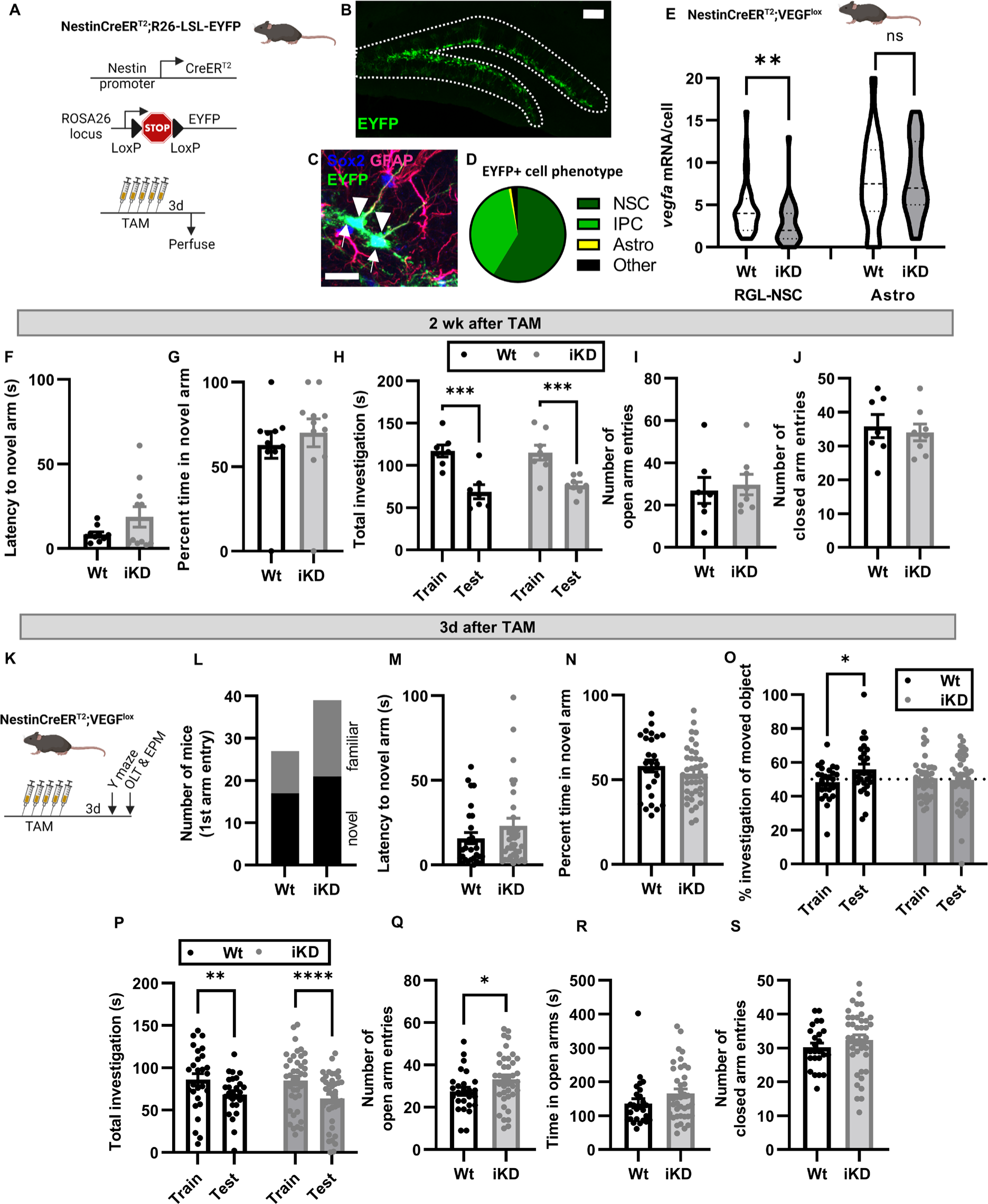
NSPC-VEGF knockdown specificity and behavioral data at 3d after TAM. A) Schematic of transgenic mouse strains used to further assess Cre-induced recombination specificity. NestinCreER^T2+/-^;Rosa-Lox-STOP-lox^+/-^ mice mice were perfused 3d after TAM. B) Representative immunofluorescent images of EYFP labeling in the DG. Scale = 100 μm. C) Representative image of EYFP+ NSCs (Sox2+/GFAP+ with apical process, arrowhead) and IPCs (Sox2+/GFAP-, arrows). Scale = 20 μm. D) Pie graph showing percent of EYFP+ cells in the DG that where phenotypic NSCs, IPCs, astrocytes or other. N = 3 mice. E) Violin plots of *vegfa* mRNA puncta per cell in Wt and iKD NSCs versus astrocytes. **p = 0.0025 Mann Whitney test. N = 40 RGL-NSCs/group, 20 astrocytes/group. F) Latency of mice to enter novel arm in Y maze. T-test, ns. N = 10 Wt, 11 iKD mice. G) Percent time in novel arm over total time in novel and familiar arm. T-test, ns. Mean ± SEM and individual mice shown. N = 10 Wt, 11 iKD. H) Total object investigation time in the object location training and testing sessions. 2-way ANOVA trial p <0.0001. ***p < 0.001 Sidak’s multiple comparisons. Mean ± SEM and individual mice shown. N = 7 Wt, 8 iKD mice. I) Number of open arm entries in the EPM. T-test, ns. Mean ± SEM and individual mice shown. N = 7 Wt, 8 iKD mice. J) Number of closed arm entries in the EPM. T-test, ns. Mean ± SEM and individual mice shown. N = 7 Wt, 8 iKD mice. K) Schematic of treatments and behavioral testing. Mice received 5d TAM then 3 days later were tested on a hippocampal dependent Y maze, hippocampal dependent object location task (OTL) and an elevated plus maze (EPM). L) Number of mice who chose the novel arm first for entry in Y maze. Fisher’s exact test, ns. M) Latency of mice to enter novel arm in Y maze. T-test, ns. Mean ± SEM and individual mice shown. N) Percent time in novel arm over total time in novel and familiar arm. T-test, ns. Mean ± SEM and individual mice shown. O) Percent time spent investigating a moved object in the OLT during training and testing. 2-way ANOVA trial x genotype interaction p = 0.0299. *p = 0.0243 Sidak’s multiple comparisons. Mean ± SEM and individual mice shown. P) Total object investigation time in the object location training and testing sessions. 2-way ANOVA trial p < 0.0001. **,***p < 0.01, 0.001 Sidak’s multiple comparisons. Mean ± SEM and individual mice shown. Q) Number of open arm entries in the EPM. *p = 0.0493, T-test. Mean ± SEM and individual mice shown. R) Time spent in open arms of the EPM. T-test, ns. Mean ± SEM and individual mice shown. S) Number of closed arm entries in the EPM. T-test, ns. Mean ± SEM and individual mice shown. L-S) N = 27 Wt, 39 iKD mice.

**Supplemental Figure 2.**
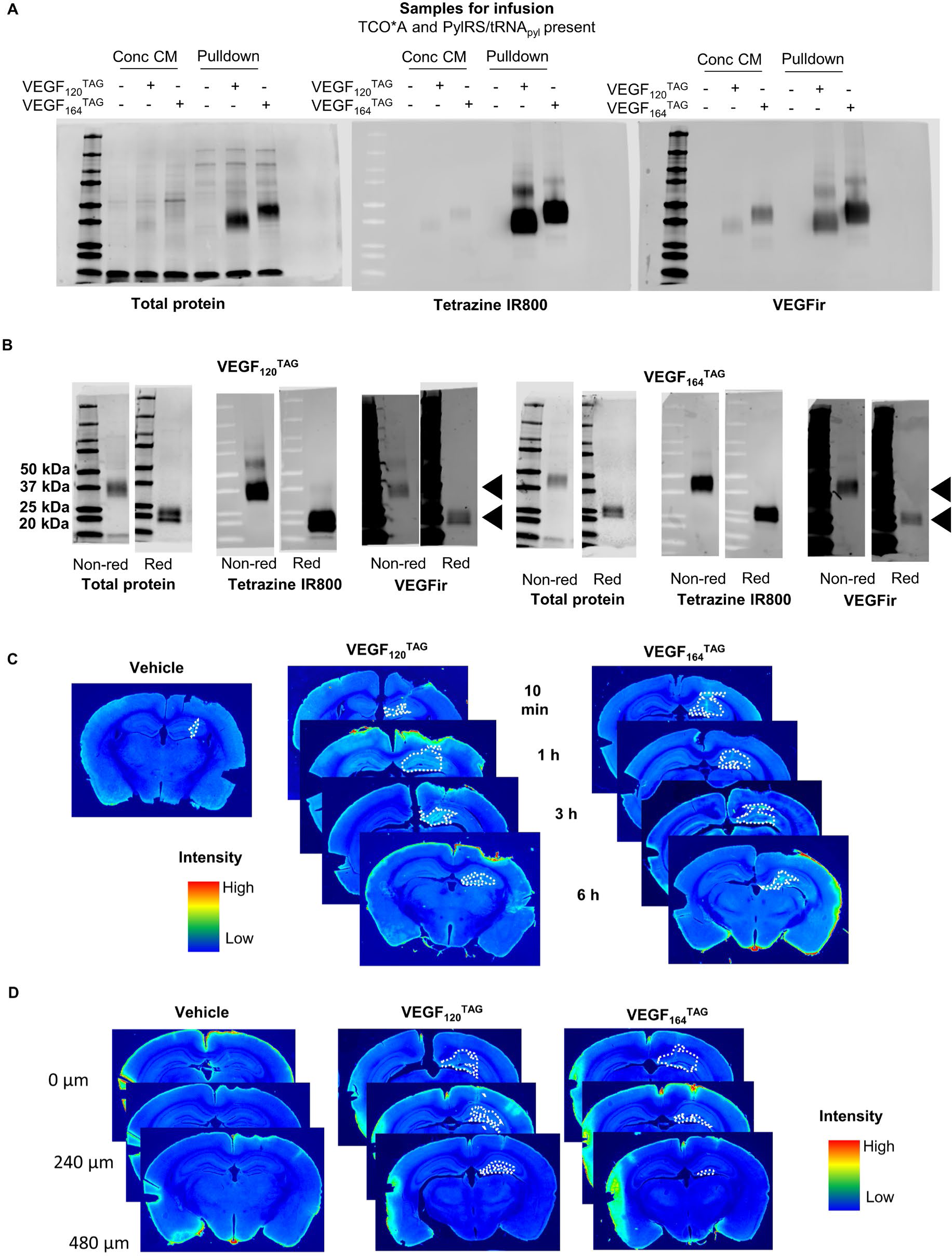
Generation and isolation of VEGF^TAG^ proteins. A) Images of western blots showing total protein, TCO*A bearing protein (reacted with tetrazine-biotin then streptavidin IR800), and VEGF immunoreactivity (ir) in whole concentrated CM and bead-concentrated final samples used for infusion. Non-reducing conditions were used, resulting in detection of VEGF dimers. B) Images of western blots run in non-reducing conditions (to reveal VEGF monomers) and reducing conditions (to reveal VEGF dimers). Arrowheads signal dimers and monomers. Blots were replicated at least twice to verify protein identity and tetrazine-reactivity. C) Representative images of VEGF^TAG^ areas over time. Dashed outline shows VEGF^TAG^+ area. Vehicle mouse, VEGF_120_^TAG^ (3h) mouse and VEGF_164_^TAG^ (1h) mouse are duplicated from main figure. D) Representative images of VEGF^TAG^ areas in single mice from epicenter to 480 µm rostral. Dashed outline shows VEGF^TAG^+ area.

**Supplemental Figure 3.**
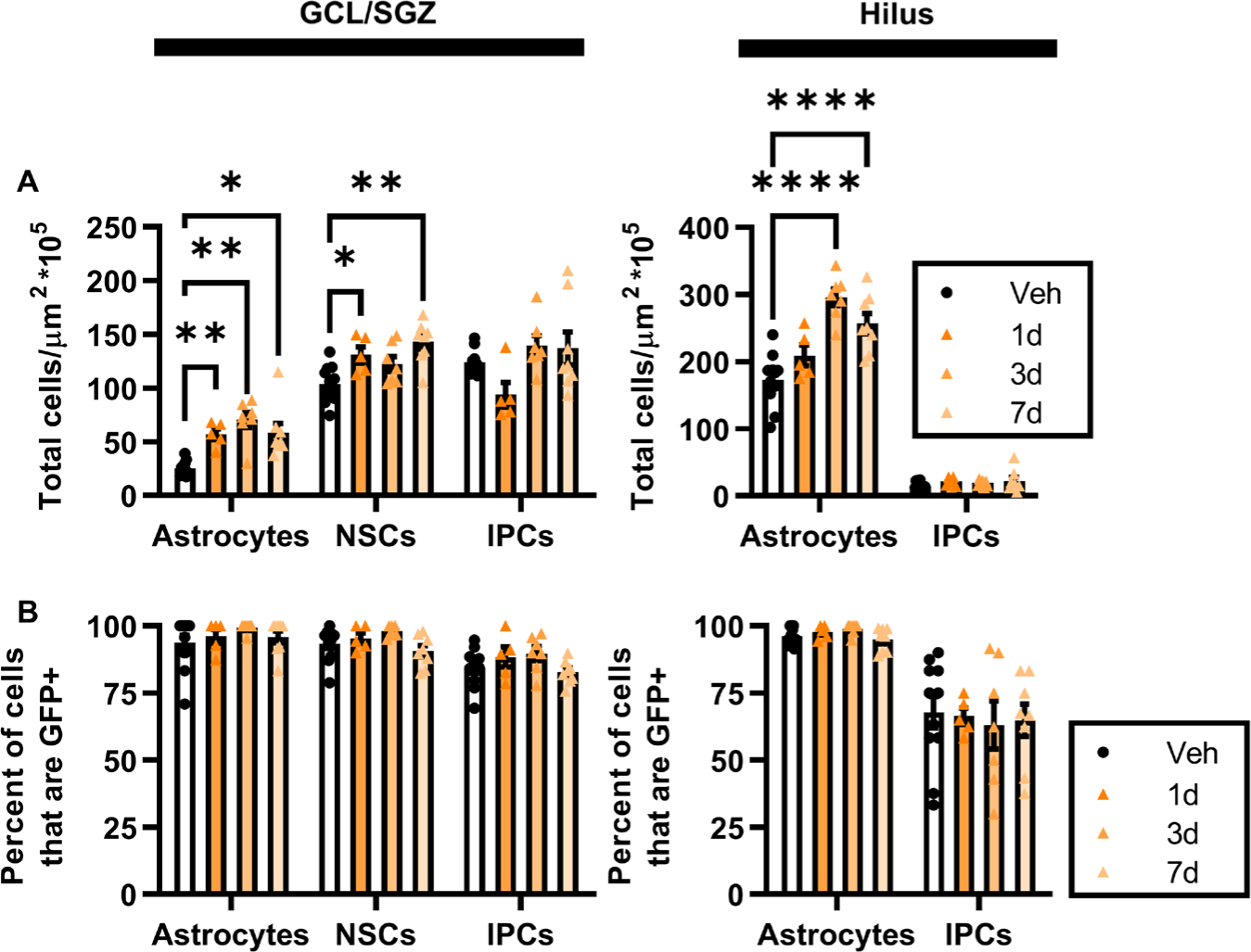
Cell proliferation response in VEGF-GFP mice after KA. A) Density of total cells in the granule cell layer/subgranular zone (GCL/SGZ) and hilus identified as astrocytes, NSCs or IPCs based on immunolabeling for Sox2 and GFAP quantified 1d, 3d, or 7d after a single systemic injection of KA in adult VEGF-GFP mice. 2 way repeated measures ANOVA: SGZ/GCL: cell type x timepoint p = 0.0027, cell type p < 0.0001, time p < 0.0001. Hilus: cell type x timepoint p < 0.0001, cell type p < 0.0001, time p < 0.0001. *,**,**** p < 0.05, 0.01, 0.0001 Dunnett’s multiple comparisons to Vehicle. B) Percent of astrocytes, NSCs and IPCs in the granule cell layer/subgranular zone (GCL/SGZ) and hilus identified that were GFP+ 1d, 3d, or 7d after a single systemic injection of KA in adult VEGF-GFP mice. 2 way repeated measures ANOVA: SGZ/GCL: cell type p < 0.0001. Hilus: cell type p < 0.0001. Dunnett’s multiple comparisons to Vehicle all ns. A,B) Mean ± SEM and individual mice shown. N = 11 vehicle, N = 5-8/KA timepoint.

**Supplemental Figure 4.**
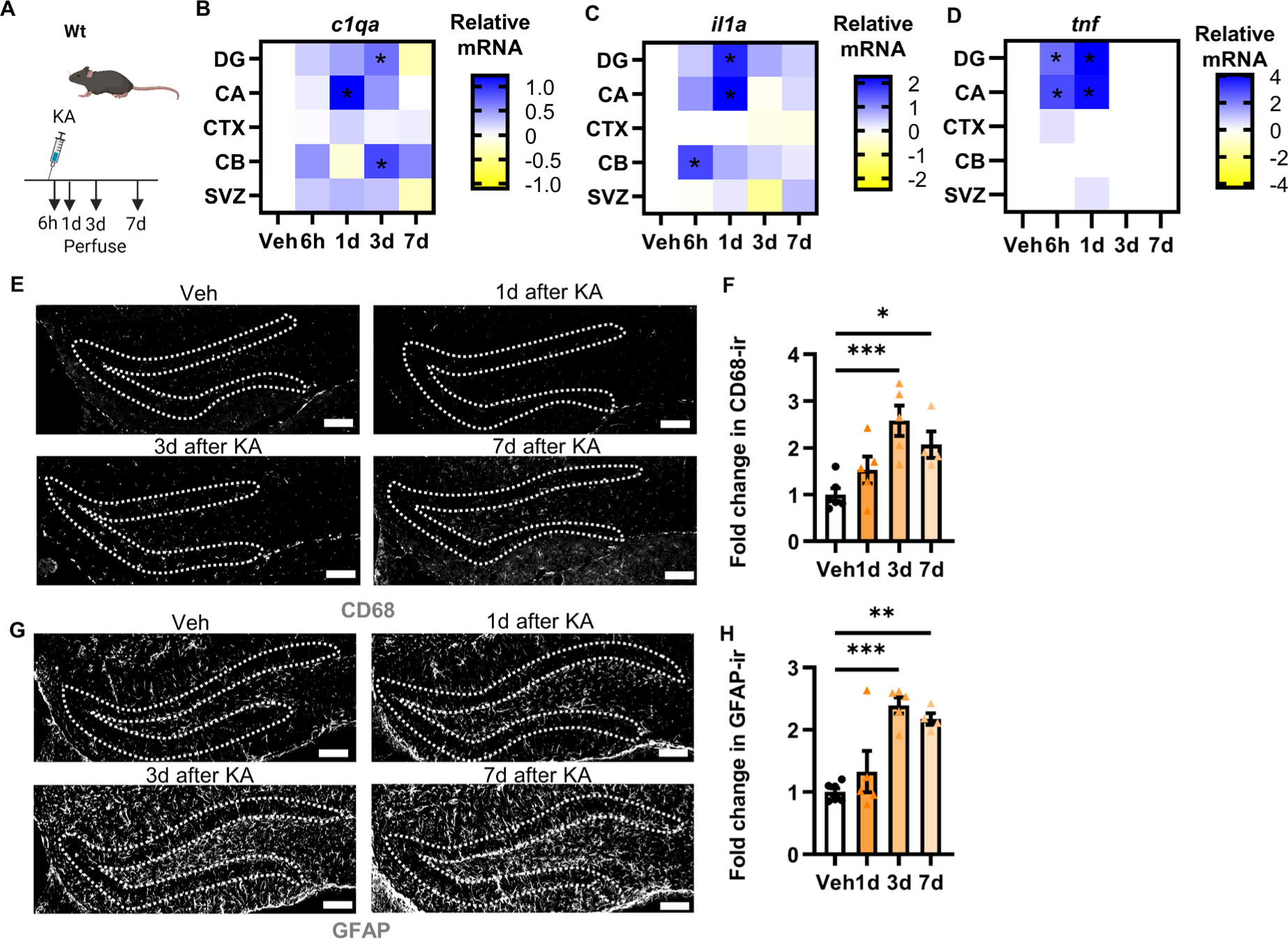
Kainic acid causes DG neuroinflammation. A) Schematic of treatments. Wt mice received a single injection of systemic KA then were perfused 6h-7d later. B-D) Relative *C1qa (B), Il1a (C)* or *Tnf (D)* mRNA levels in dentate gyrus (DG), CA regions of hippocampus (CA), cortex (CTX), cerebellum (CB) and subventricular zone (SVZ) after KA injection determined by real time qPCR. Two-way ANOVA: C1qa area x treatment interaction p = 0.0009, treatment p < 0.0001. Il1a area x treatment interaction p = 0.0005, treatment p < 0.0001, area p < 0.0001. Tnf area x treatment interaction p < 0.0001, treatment p < 0.0001, area p < 0.0001. *p < 0.05 Dunnett’s multiple comparisons versus vehicle within area. N = 20-23 Veh mice, 6-12 KA treated mice/area/timepoint. E) Representative image of CD68 immunoreactivity (ir) in the DG of Wt mice 1d after KA injection. Scale = 100 µm. F) Fold change in CD68 immunoreactivity (ir) thresholded area in the DG of Wt mice 1-7d after KA relative to Veh mice. ANOVA p =0.0023, *,***p < 0.05, 0.001 Dunnett’s multiple comparisons to Veh. Mean ± SEM and individual mice shown. N = 4-6 mice/group. G) Representative image of GFAP immunoreactivity (ir) in the DG of Wt mice 1-7d after KA injection. Scale = 100 µm. H) Fold change in GFAP immunoreactivity (ir) thresholded area in the DG of Wt mice 1-7d after KA relative to Veh treated mice. ANOVA p = 0.0001, **,***p < 0.01, 0.001 Dunnett’s multiple comparisons to Veh. Mean ± SEM and individual mice shown. N = 4-6 mice/group.

**Supplemental Figure 5.**
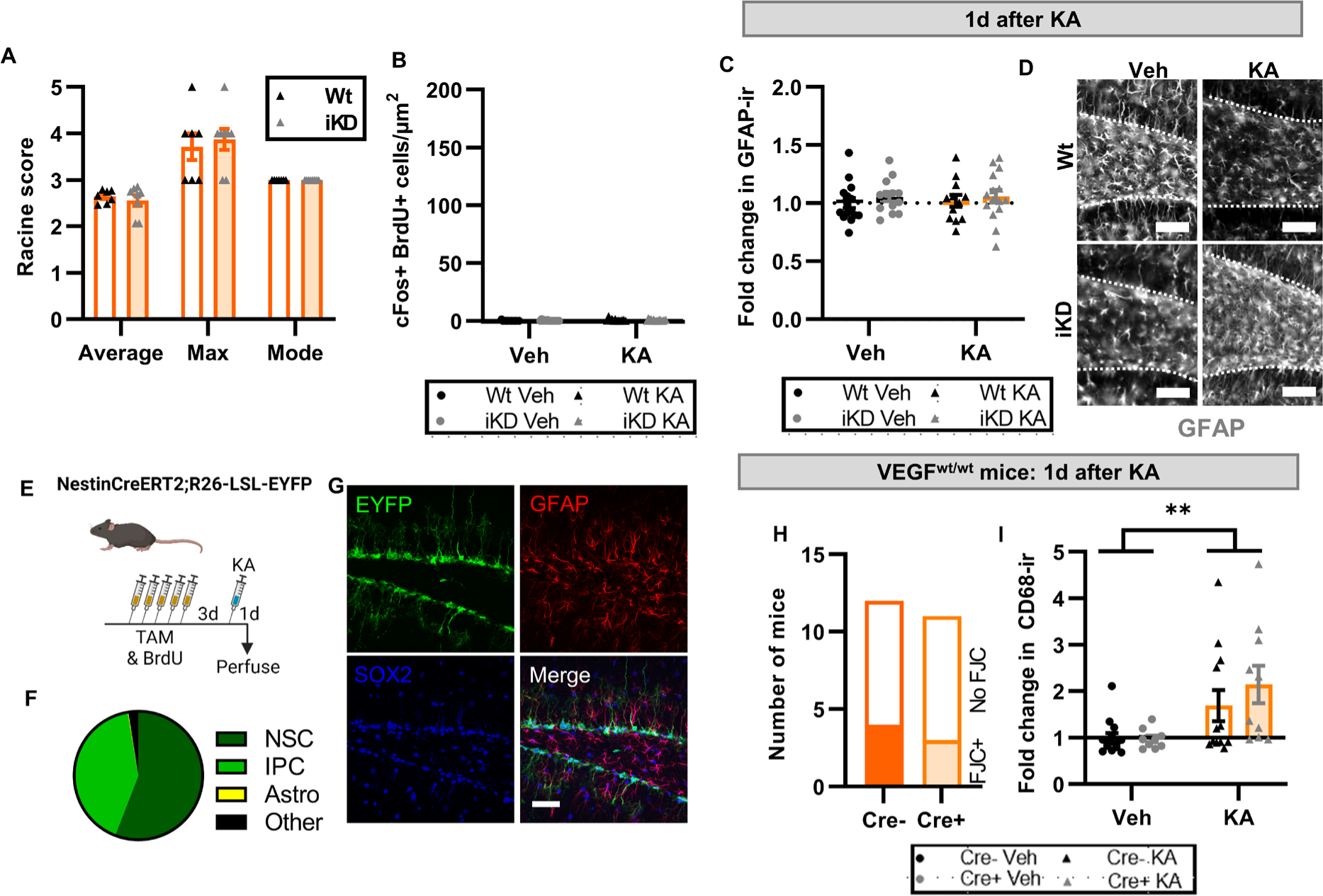
NSPC-VEGF knockdown exacerbates excitotoxic injury. A) Average, maximum and mode seizure score of Wt and iKD mice according to the Racine scale in the 4h following KA injection. T-tests within each category all ns. Mean ± SEM and individual mice shown. N = 7-8 mice/genotype. B) Density of BrdU+cFos+ cells in the granule cell layer 1d after KA injection in Wt and iKD mice. 2 way ANOVA treatment p < 0.0001. C) Fold change in GFAP immunoreactivity (ir) thresholded area in the DG of Wt and iKD mice 1d after KA relative to Wt-Veh mice. 2 way ANOVA all ns. Mean ± SEM and individual mice shown. N = 12-15 mice/group. D) Representative images of GFAPir in the DG of Wt and iKD mice. Scale = 20 µm. E) Schematic of treatments. NestinCreER^T2+/-^;LoxP-STOP-LoxP-EYFP^+/-^ mice received 5d TAM injections followed by a single KA injection 3d after the last TAM injection. Mice were perfused 1d after KA. F) Pie graph showing percent of EYFP+ cells in the DG that where phenotypic NSCs, IPCs, astrocytes or other 1d after KA. N = 3 mice. G) Representative image of EYFP col-labeling with Sox2 and GFAP used to identify astrocytes, NSCs and IPCs. Scale = 20 µm. H) Number of NestinCreER^T2−/−^ and NestinCreER^T2+/-^ mice showing any FJC labeling in the DG 1 d after KA. Fisher’s exact test ns. I) Fold change in CD68 immunoreactivity (ir) thresholded area in the DG of NestinCreER^T2−/−^ and NestinCreER^T2+/-^ mice 1d after KA relative to NestinCreER^T2−/−^ Veh treated mice. 2 way ANOVA treatment p = 0.0011. Mean ± SEM and individual mice shown. N = 9-14 mice/group.

